# Factor cooperation for chromosome discrimination in *Drosophila*

**DOI:** 10.1101/414193

**Authors:** Christian Albig, Evgeniya Tikhonova, Silke Krause, Oksana Maksimenko, Catherine Regnard, Peter B. Becker

## Abstract

Transcription regulators select their genomic binding sites from a large pool of similar, non-functional sequences. Although general principles that allow such discrimination are known, the complexity of DNA elements often precludes a prediction of functional sites.

The process of dosage compensation in *Drosophila* allows exploring the rules underlying binding site selectivity. The male-specific-lethal (MSL) Dosage Compensation Complex selectively binds to some 300 X-chromosomal ‘High Affinity Sites’ (HAS) containing GA-rich ‘MSL recognition elements’ (MREs), but disregards thousands of other MRE sequences in the genome. The DNA-binding subunit MSL2 alone identifies a subset of MREs, but fails to recognize most MREs within HAS. The ‘Chromatin-linked adaptor for MSL proteins’ (CLAMP) also interacts with many MREs genome-wide and promotes DCC binding to HAS. Using genome-wide DNA-immunoprecipitation we describe extensive cooperativity between both factors, depending on the nature of the binding sites. These are explained by physical interaction between MSL2 and CLAMP. *In vivo*, both factors cooperate to compete with nucleosome formation at HAS. The male-specific MSL2 thus synergises with a ubiquitous GA-repeat binding protein for refined X/autosome discrimination.

## Introduction

The rules according to which transcription factors select only a small subset of genomic binding sites from a large excess of similar sequences remain largely elusive. Typically, the binding sites for transcription factors consist of short sequence motifs, yet only a few percent of all genomic sites that conform to a consensus motif are functional and bound by protein *in vivo*. This selectivity may be explained by cooperativity between factors binding complex DNA elements (Jolma et al., 2015; Mirny, 2010), the instructive role of DNA conformation (Abe et al., 2015; Dror et al., 2015; Slattery et al., 2014; Watson et al., 2013) and sequence context as well as the role of chromatin organization, which may occlude non-functional sites (Li et al., 2011; Soufi et al., 2015).

A striking example is the process of sex chromosome dosage compensation in *Drosophila melanogaster*, which doubles the transcription output of most genes on the single X chromosome in males to match the two active X chromosomes in females. This sophisticated regulation is brought about by the male-specific-lethal dosage compensation complex (MSL-DCC, or just DCC). The DCC consists of five ‘male-specific-lethal’ (MSL) protein subunits (MSL1, MSL2, MSL3, MOF [males-absent-on-the-first] and MLE [maleless]) and two long, non-coding *roX* RNAs (RNA-on-the-X) [reviewed in (Keller and Akhtar, 2015; Kuroda et al., 2016; Lucchesi and Kuroda, 2015)].

Dosage compensation is genetically encoded on the X chromosome in the form of ∽300 high affinity binding sites (HAS) for the DCC, which are also referred to as ‘chromosomal entry sites’ (CES). The current model poses that the DCC first interacts with HAS on the X chromosome and then transfers to active genes in its vicinity [(Schauer et al., 2017) and reviewed in (Kuroda et al., 2016; Samata and Akhtar, 2018)]. These genes are epigenetically marked by methylation of histone H3 at lysine 36 (H3K36me3), a mark that is placed co-transcriptionally. The DCC subunit MSL3 contains a chromo-barrel domain that serves as a ‘reader head’ to scan the chromatin for the active methylation mark (Larschan et al., 2007; Prestel et al., 2010). Upon binding, the associated ‘writer’ subunit MOF acetylates histone H4 at lysine 16 (H4K16) (Akhtar and Becker, 2000; Gelbart et al., 2009; Smith et al., 2000), which somehow boosts the production of functional mRNA through unfolding of the chromatin fiber (Ferrari et al., 2014). Any gene integrated on the X chromosome is subject to this regulation. Understanding dosage compensation, therefore, requires understanding the nature of X-specific DCC binding.

The HAS harbor a low-complexity, GA-rich consensus motif, referred to as ‘MSL recognition element’ (MRE) (Alekseyenko et al., 2008; Straub et al., 2008), which is indispensable for DCC binding. However, the genome contains several thousand MREs on the X chromosome outside of HAS and on autosomes, therefore only ∽2% of MREs are functional and bound by the DCC (Alekseyenko et al., 2008; Straub et al., 2008). The direct MSL2 binding sites have been experimentally determined by *in vitro* genome-wide DNA immunoprecipitation assays (Villa et al., 2016). MSL2 binds to DNA via a C-terminal CXC domain followed by a region rich in prolines (Fauth et al., 2010; Zheng et al., 2014). Remarkably, the CXC domain recognizes a subset of MREs whose consensus motif has a notable 5’ extension characterized by a particular DNA shape (Villa et al., 2016). These CXC-dependent sites are named ‘Pioneering-sites-on-the-X’ (PionX), as they (1) are the first to be bound upon *de novo* induction of dosage compensation in females, (2) are preferentially contacted by an MSL2-MSL1 sub-complex, and (3) are enriched on the evolutionary young neo-X chromosome of *Drosophila miranda* (Dahlsveen et al., 2006; Villa et al., 2016). The PionX motif is superior over the MRE motif in predicting which genomic sites function as HAS. The PionX motif is up to ∽10-fold enriched on the X chromosome, providing a first clue about how MSL2 distinguishes the X chromosome from autosomes (Villa et al., 2016). In general, however, the interaction of MSL2 with PionX sites does not fully explain HAS targeting, since only a small fraction of the MSL2 *in vitro* binding sites (mostly containing a PionX signature) overlap with functional HAS *in vivo*.

A solution to the problem was suggested by Larschan and colleagues, who found a zinc finger protein that associates with about half of MREs throughout the genome. They termed this protein CLAMP (Chromatin Linked Adaptor for MSL Proteins) since depletion of the protein leads to dissociation of the DCC from the X chromosome (Larschan et al., 2012; Soruco et al., 2013). CLAMP is an essential protein in *Drosophila* independent of sex, which binds thousands of GA-rich sequences genome-wide (Kuzu et al., 2016; Soruco et al., 2013; Urban et al., 2017b) and therefore does not qualify as a determinant of X-specificity. Remarkably, CLAMP binds to HAS only in male cells, suggesting a functional relationship with the DCC (Soruco et al., 2013).

It is possible that CLAMP facilitates MSL2 binding to MREs by keeping these elements nucleosome-free, in analogy to early observations that the GAGA factor (GAF) keeps promoters and polycomb response elements clear of nucleosomes to allow other regulators to bind (Leibovitch et al., 2002; Lu et al., 1993; Strutt et al., 1997; Tsukiyama et al., 1994). Indeed, Urban et al. recently found that CLAMP promotes the accessibility of DNA in chromatin over long distances surrounding its binding sites (Urban et al., 2017a). This Micrococcus Nuclease (MNase) titration study also suggested that CLAMP leads to a global decompaction of the X chromosome in males.

To explore the relationship between CLAMP and MSL2 we integrated data from several approaches. We monitored how the two factors influenced each other’s binding to genomic sequences *in vitro* by DNA immunoprecipitation (Gossett and Lieb, 2008; Liu et al., 2005; Villa et al., 2016). We observed mutual recruitment, explained by direct interaction between both proteins and shared affinity for long GA-repeat sequences. This DNA binding cooperativity improved reliable selection of functional MREs which are located within HAS, however at the expense of binding to additional, non-functional sites. To explore whether the chromatin organization of the genome plays a role, we monitored DNA accessibility genome-wide in S2 and Kc cells by ATAC-seq (Assay for Transposase Accessibly Chromatin with high-throughput sequencing) (Buenrostro et al., 2013; Buenrostro et al., 2015) and observed how the pattern of nucleosome-free chromatin changed upon RNA interference with CLAMP or MSL2 expression. We integrated these data with direct measurements of the *in vivo* interactions of both proteins by ChIP-seq (Chromatin Immunoprecipitation with high-throughput sequencing) (Straub et al., 2013). The data do not support the hypothesis of a hierarchical relationship between the two proteins. Rather, both factors synergize to keep common binding sites nucleosome-free and stabilize each other’s binding. We conclude that the correct targeting of X chromosomal HAS involves synergistic action of the male-specific MSL2 and the general GA-repeat binding CLAMP to compete with nucleosome formation.

## Results

### MSL2 requires CLAMP for efficient binding to HAS *in vivo*

Larschan and co-worker previously reported that binding of MSL3 to the polytenic X chromosome and the crosslinking of MSL2 to some HAS required the presence of the CLAMP protein, which recognizes GA-rich sequences matching the MRE motif (Larschan et al., 2012; Soruco et al., 2013). On the other hand, recombinant MSL2, the male-specific DNA-binding subunit of the DCC, can select MRE sequences and is able to enrich for X chromosomal sites, especially those with PionX signature, in genomic DNA *in vitro* (Villa et al., 2016).

To explore the relationship between CLAMP and MSL2 in our system, we depleted CLAMP by RNA interference (RNAi) in male S2 cells and monitored the X chromosome binding of MSL3 and MSL2 by immunostaining. RNAi against *clamp* or *msl2* is efficient and does not affect each other’s levels, excluding indirect effects (Fig. 1A). An irrelevant RNAi against green fluorescent protein (*gfp*) or *Schistosoma japonicum* glutathione-S-transferase (*gst*) served as control (‘control’ in all figures). Upon *clamp* RNAi, MSL2 and MSL3 no longer localize at a coherent X chromosome territory, but redistribute in smaller speckles, indicating a targeting deficiency (Fig. 1B). The fact that both subunits still co-localize suggests that the DCC is still intact and rather mis-targeted as a complex.

**Figure 1.**
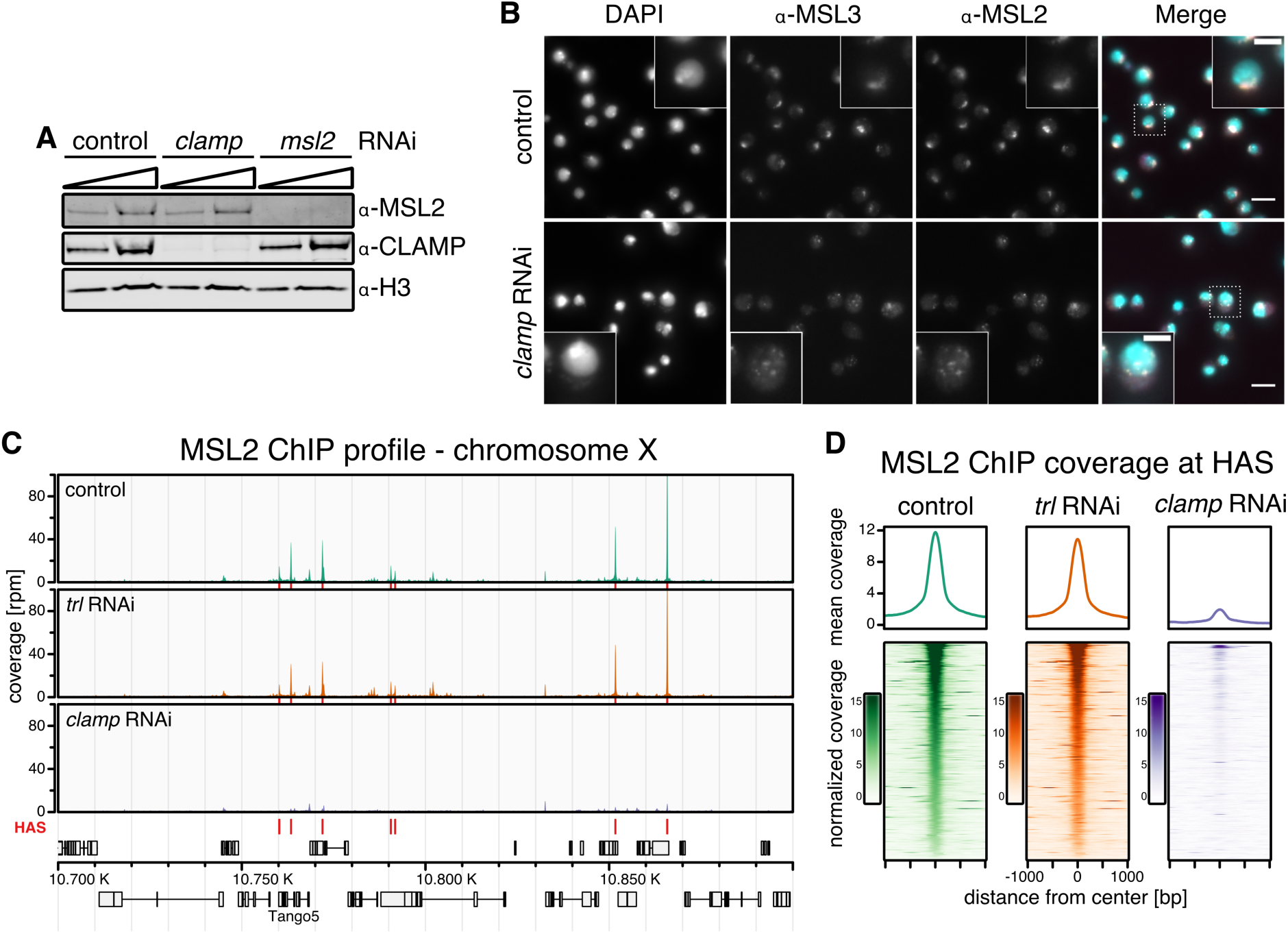
Binding of MSL2 to HAS in male S2 cells depends on CLAMP. **(A)** Western blot detection of MSL2, CLAMP and H3 in whole cell extracts from S2 cells after *msl2* or *clamp* RNAi. An irrelevant RNAi directed against green fluorescent protein (*gfp*) or *Schistosoma japonicum* glutathione-S-transferase (*gst*) sequences serve as control for these and further experiments. **(B)** Immunofluorescence microscopy of MSL2 and MSL3 in control cells and upon *clamp* RNAi. The merged images show MSL3 in magenta, MSL2 in yellow and DAPI staining in cyan. A region of zoom-in is marked by dashed rectangle. Scale bar: 10 µm (5 µm in inset). **(C)** Genome browser profile of MSL2 ChIP seq profile showing mean coverage (control cells n = 3, *trl* RNAi: n = 3, *clamp* RNAi: n = 4) along a representative 200 kb window on the X chromosome. Red bars above the gene models indicate location of HAS. **(D)** Average profile and heat map of mean MSL2 ChIP seq coverage (control cells n = 3, *trl* RNAi: n = 3, *clamp* RNAi: n = 4) in a 2 kb window centered on 309 HAS as indicated. Heat maps are sorted by decreasing MSL2 enrichment in peak region in the control data.

To further investigate MSL2 binding *in vivo* in absence of CLAMP, we performed ChIP-seq of MSL2 in male S2 cells after depletion of CLAMP by RNAi (Fig. EV1A). Upon depletion, the CLAMP ChIP-seq signal was robustly reduced and only minor residual binding was observable (Fig. EV1B). Under those conditions MSL2 binding to HAS was massively reduced (Figs. 1C and D). HAS with a PionX signature are the primary contacts for MSL2 *in vivo* (Villa et al., 2016). MSL2 binding to the subset of HAS which overlap with PionX sites (HAS-PionX) seemed to be slightly less affected by *clamp* RNAi, however the small amounts of remaining CLAMP after RNAi also tended to be enriched at HAS-PionX sites (Figs. EV1C and D). The dependence of MSL2 on CLAMP for binding to all HAS *in vivo* was somewhat unexpected as MSL2 can target to PionX sites without the help of any additional factor *in vitro*.

As a control for these experiments we depleted the GAGA factor (GAF). GAF (encoded by the *trithorax-like [trl]* gene) also binds to GA-rich sites genome-wide and co-localizes with CLAMP at many sites (Soruco et al., 2013). Contrasting CLAMP, GAF is absent from most HAS (Alekseyenko et al., 2012; Greenberg et al., 2004; Straub et al., 2008). As expected, the depletion of GAF did not affect MSL2 binding (Figs 1C, D).

Since Larschan and colleagues suggested some competition in DNA-binding between CLAMP and GAF (Kaye et al., 2018), we explored whether the two proteins compete for binding to HAS *in vivo*. Upon RNAi depletion of GAF in male S2 cells the CLAMP ChIP-seq signal was unchanged at HAS (Figs. EV1B, D). Genome-wide, at a false discovery rate (FDR) < 0.05, only 93 out of 5983 CLAMP binding sites (1.55%) showed a statistically significant difference in CLAMP binding (Fig. EV1E). These sites were mostly located in chromatin with enhancer- or promoter-related histone marks and CLAMP binding was increased or decreased (Appendix S1).

To analyze potential competition between GAF and CLAMP, we also monitored GAF binding at CLAMP binding sites after depletion of CLAMP (Fig. EV1F). Nine robust CLAMP binding sites were monitored by ChIP-qPCR including three HAS (Fig. EV1G). At none of these sites did GAF binding change upon removal of CLAMP.

We conclude that MSL2/DCC binding to HAS depends on CLAMP but not on GAF and that CLAMP and GAF do not generally compete for binding to GA-rich sequences.

### Genomic binding sites of CLAMP defined *in vitro* by DNA immunoprecipitation

We previously assayed the DNA binding sites of MSL2 *in vitro* through genome-wide DNA immunoprecipitation (DIP) (Gossett and Lieb, 2008; Liu et al., 2005; Villa et al., 2016). In such experiments, purified and fragmented genomic DNA is incubated with protein of interest under conditions where competitive DNA binding occurs. The protein is immunoprecipitated and the bound DNA sequenced. For CLAMP, such an analysis had not been done. We expressed and purified recombinant full-length CLAMP via a baculovirus expression system (Appendix S2) and assayed it’s *in vitro* DNA-binding property by genome-wide DIP-seq.

*In vitro*, CLAMP bound to 4037 sites in the *Drosophila* genome under our DIP conditions (Fig. 2A), comparable in number to 5214 sites determined by our ChIP-seq approach. About one third (32.3%, n = 1307) of the *in vitro* CLAMP binding sites overlapped with *in vivo* binding sites (Fig. 2B). Sequence analyses within *in vitro* binding sites yielded the core of the GA-repeat consensus motif defined *in vivo* by ChIP-seq, replicating the previously identified CLAMP motif, which has high similarity to the MRE motif (Fig. 2C) (Soruco et al., 2013). Furthermore, the CLAMP *in vitro* binding motif is highly similar to the previously identified motif for the CLAMP DNA-binding domain *in vitro* by protein-binding microarray, confirming the DNA binding specificity for our full-length recombinant protein (Kuzu et al., 2016; Soruco et al., 2013). As expected, CLAMP bound sites on all chromosomes *in vitro* with a 2.4-fold enrichment of X chromosomal sequences similarly to *in vivo* with 1.7-fold enrichment (Fig. 2D), reflecting the ∽2-fold enrichment of the MRE motif on the X chromosome (Alekseyenko et al., 2008; Straub et al., 2008). However, the binding intensities *in vitro* and *in vivo* were uncorrelated (Fig. 2E). Whereas the *in vitro* binding sites solely reflect the intrinsic binding affinity of CLAMP to DNA, the *in vivo* interactions are obviously modulated by other factors (such as MSL2, see below).

**Figure 2.**
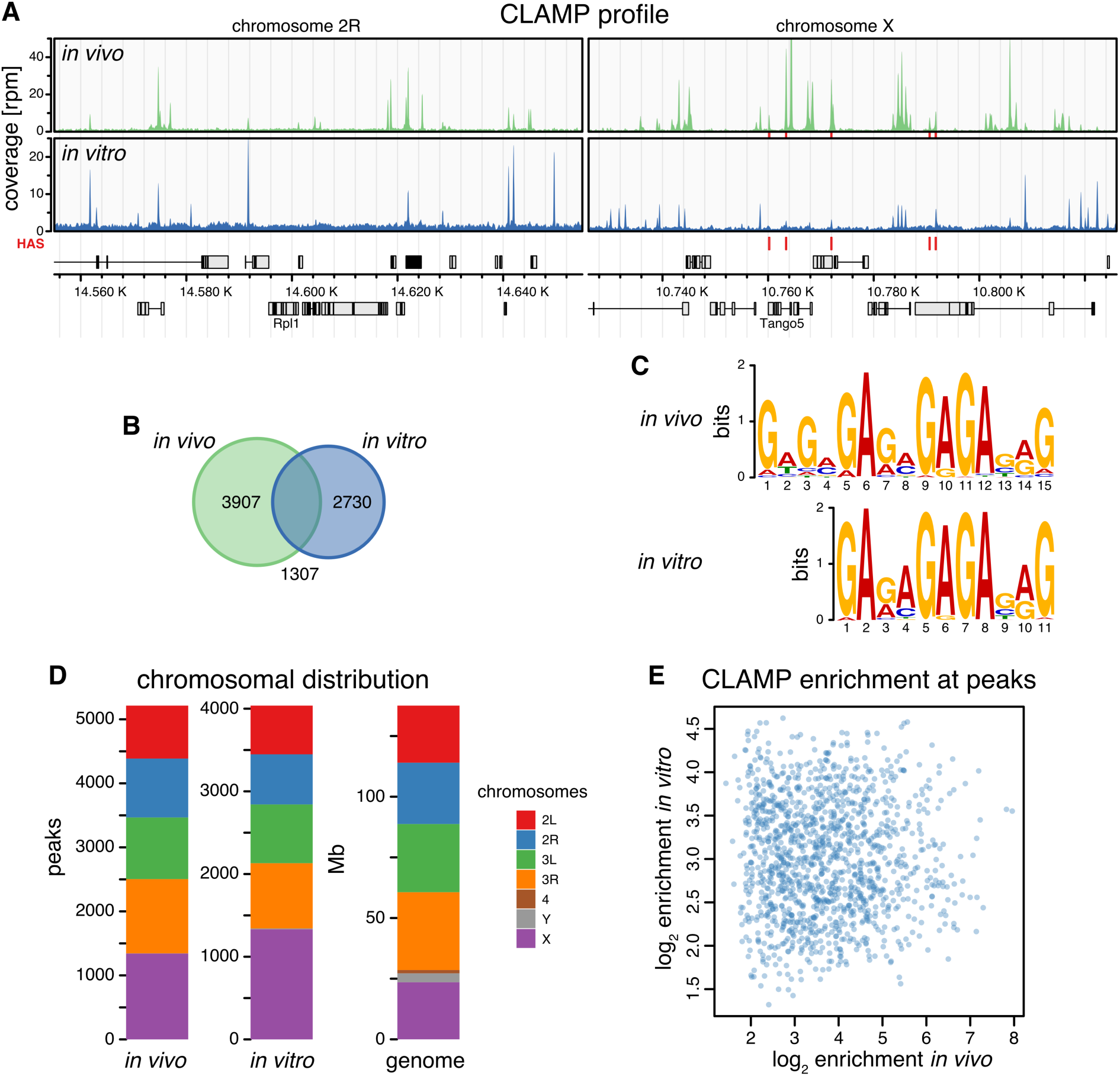
CLAMP selects GA-rich consensus sequence motifs *in vitro*. **(A)** Genome browser profile of *in vivo* CLAMP ChIP-seq (upper panel) and *in vitro* CLAMP DIP-seq (lower panel) profiles (mean coverages, n = 3 each) along representative 200 kb windows on chromosome 2R and X. Red bars above the gene models indicate positions of HAS. **(B)** Venn diagram of robust peak sets from *in vivo* CLAMP ChIP-seq (n = 5214) and *in vitro* CLAMP DIP-seq (n = 4037). **(C)** *De novo* discovered motifs from robust peak sets as in (B). **(D)** Bar chart of chromosomal distribution of robust peak sets as in (B). The chromosome sizes serve as reference for uniform distribution (genome). **(E)** Scatterplot of *in vitro* CLAMP DIP-seq mean log_2_ enrichment (n = 3) against *in vivo* CLAMP ChIP-seq mean log_2_ enrichment (n = 3) at 1307 overlapping peak regions displayed on (B).

### Cooperation between CLAMP and MSL2 increases the selectivity of HAS recognition in the context of the genome

We recently described the *in vitro* genomic binding sites of MSL2 by DIP-seq (Villa et al., 2016). This study revealed the capacity of MSL2 to identify PionX sites and to enrich X-chromosomal sequences, but the protein missed many HAS and in addition pulled out autosomal GA-rich sites that do not correspond to physiological binding sites (Villa et al., 2016). We now explored whether cooperation with CLAMP may improve the binding specificity and/or capacity of MSL2. Therefore, we expressed and purified recombinant full-length CLAMP and MSL2 via a baculovirus expression system (Appendix S2) and assayed their DNA binding properties by genome-wide DIP-seq.

In our previous study, MSL2 protein was immunoprecipitated with an *α*-FLAG antibody recognizing the epitope of its FLAG-tag. Because CLAMP protein is also FLAG-tagged to facilitate its purification, we could not use the *α*-FLAG antibody, but performed the DIP experiments using *α*-CLAMP and *α*-MSL2 antibodies specific for the proteins. The analysis is complicated by the fact that the *α*-MSL2 antibody available to us yielded a DIP profile for MSL2 without significant enrichments of sites under our conditions (Fig. 3A). Remarkably however, the *α*-MSL2 antibody retrieved a robust profile for MSL2 in the presence of CLAMP. For reference, we included our previously published MSL2 DIP-seq data using the *α*-FLAG antibody for immunoprecipitation as proxy for MSL2 binding *in vitro* (Villa et al., 2016).

**Figure 3.**
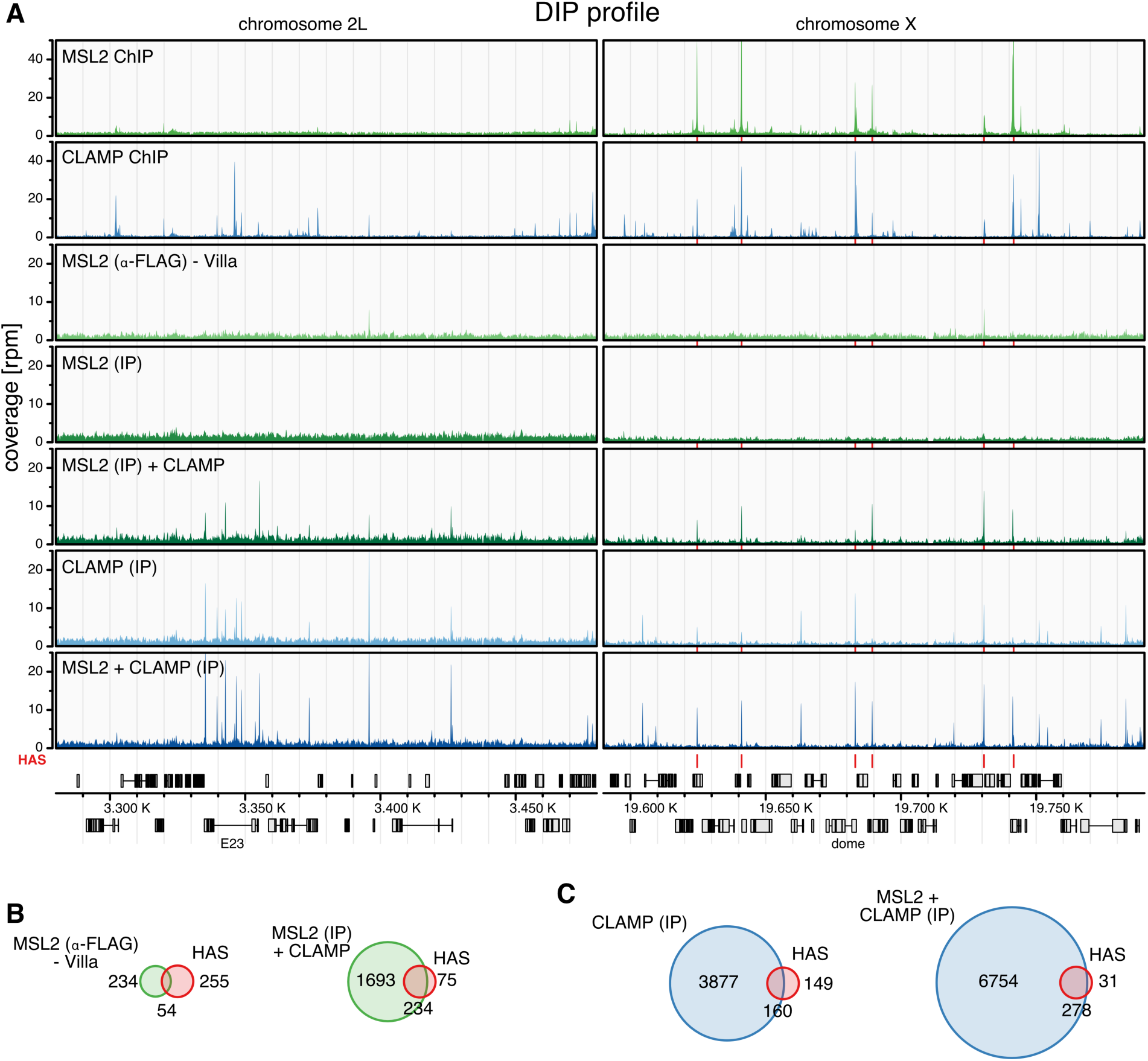
Intrinsic DNA binding cooperativity between CLAMP and MSL2 *in vitro*. **(A)** Genome browser profile showing genomic MSL2 and CLAMP binding profiles as indicated. The MSL2 and CLAMP *in vivo* ChIP-seq (top 2 profiles) and *in vitro* DIP-seq profiles (bottom 5 profiles) represent mean coverages (n = 3) along representative 200 kb windows on chromosome 2L and X. *In vitro* DIP-seq panels depict the following conditions from top to bottom: MSL2 with *α*-FLAG IP from Villa et al.(Villa et al., 2016) as proxy for MSL2 *in vitro* binding, MSL2 with *α*-MSL2 IP, MSL2 plus CLAMP with *α*-MSL2 IP, CLAMP with *α*-CLAMP IP, and MSL2 plus CLAMP with *α*-CLAMP IP. Red bars above the gene model and between the panels indicate positions of HAS. **(B)** Venn diagrams relating robust peak sets from *in vitro* DIP-seq (green) to HAS (red, n = 309). Left panel: MSL2 with *α*-FLAG IP from Villa et al.(Villa et al., 2016) as proxy for MSL2 *in vitro* binding (n = 288, overlapping n = 54). Right panel: MSL2 plus CLAMP with *α*-MSL2 IP (n = 1972, overlapping n = 234). **(C)** Venn diagrams relating robust peak sets from *in vitro* DIP-seq (blue) to HAS (red, n = 309). Left panel: CLAMP with *α*-CLAMP IP (n = 4037, overlapping n = 160). Right panel: MSL2 plus CLAMP with *α*-CLAMP IP (n = 7032, overlapping n = 278).

MSL2 alone binds *in vitro* to 288 sites of which 54 are HAS (out of 309 HAS in total), as reported previously (Villa et al., 2016) (Fig. 3B). In the presence of CLAMP, MSL2 bound now to 1927 sites including 234 HAS. A similar situation was observable for CLAMP binding *in vitro*. CLAMP alone bound 4037 sites including 160 HAS. In presence of MSL2, CLAMP bound 7032 sites including 278 HAS (Fig. 3C). This reveals an intrinsic cooperativity between MSL2 and CLAMP through stabilizing each other’s interactions at many common binding sites in the genome under these conditions. This cooperativity allows both factors together to select most HAS from genomic DNA *in vitro*, but at the cost of an increased number of non-physiological binding sites, illustrating the intrinsic binding properties of both proteins (see below, Appendix S3).

To explore whether different modes of binding cooperativity exist we performed hierarchical clustering of the *in vitro* binding intensities at the combined robust peak set of all DIP-seq samples (Fig. 4). We identified 12 major clusters defined by the different binding behaviors of both proteins. We categorized the clusters into four binding scenarios (Figs. 5A, B, EV2 and Appendix S4A): (1) independent: CLAMP and MSL2 bind alone and the presence of the other factor makes little difference [clusters 10 and 11]; (2) CLAMP-dependent: CLAMP binds largely independently of MSL2 and recruits MSL2 [clusters 6-9 (clusters 2-4 show similar behavior with lower signal enrichment)]; (3) MSL2-dependent: MSL2 binds largely independently of CLAMP and recruits CLAMP [cluster 12]; (4) interdependent: CLAMP and MSL2 do not bind alone, but show cooperative binding [cluster 5 (cluster 1 shows similar behavior with lower signal enrichment)]. As expected, MSL2 binding to PionX sites did not depend on CLAMP (Fig. 4 and 5A). CLAMP alone is absent from many PionX sites and is recruited by MSL2 to these sites.

**Figure 4.**
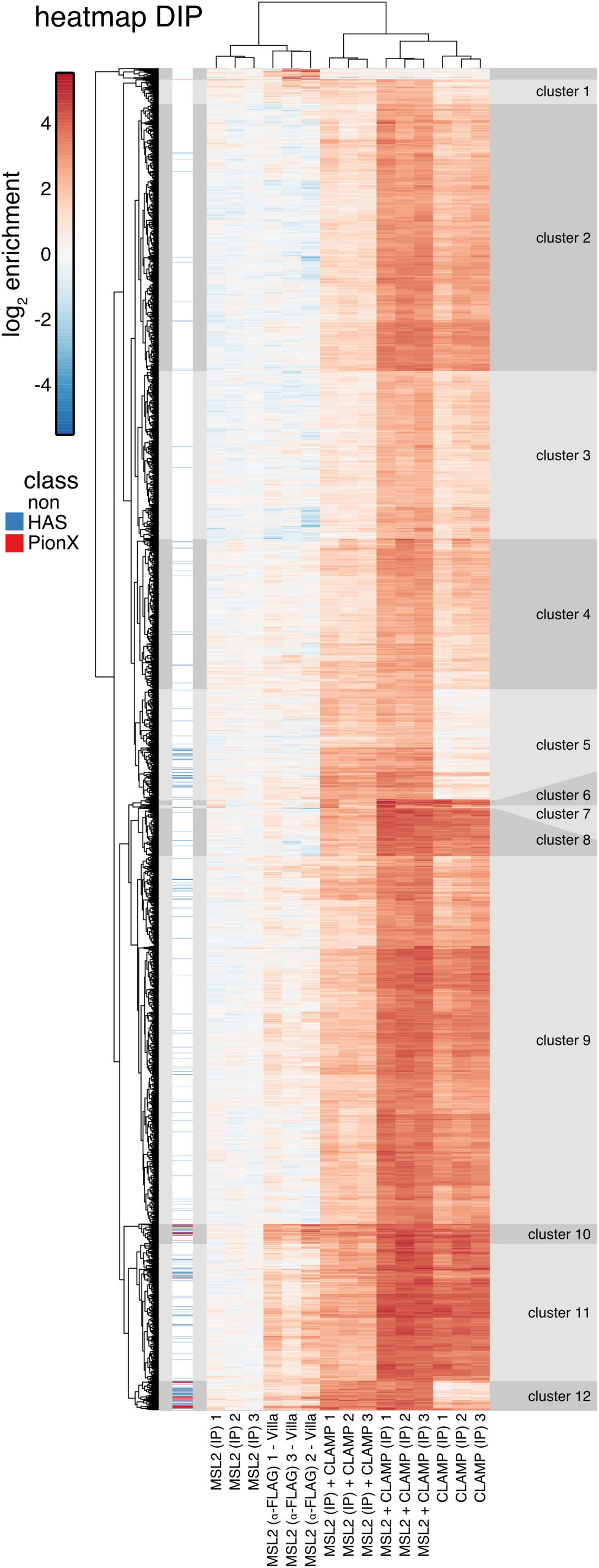
Genome-wide DNA binding of CLAMP and MSL2 *in vitro*. Clustered heat map of *in vitro* DIP-seq signal enrichment from reactions containing either MSL2 or CLAMP alone, or both proteins, as follows - the target of immunoprecipitation (IP) is indicated in brackets: MSL2 (IP *α*-FLAG) from Villa et al.,(Villa et al., 2016) as proxy for MSL2 *in vitro* binding; MSL2 (IP *α*-MSL2), MSL2 and CLAMP (IP *α*-MSL2); CLAMP (IP *α*-CLAMP); MSL2 and CLAMP (IP *α*-CLAMP) at all combined robust peaks (n = 7119). For each reaction three independent replicates are shown. Hierarchical clustering revealed 18 clusters. Clusters 1 – 12 had distinct MSL2 and CLAMP binding properties, the remaining 6 clusters at top of the heat map are small and show inconsistent enrichment between MSL2 with *α*-FLAG IP replicates.

**Figure 5.**
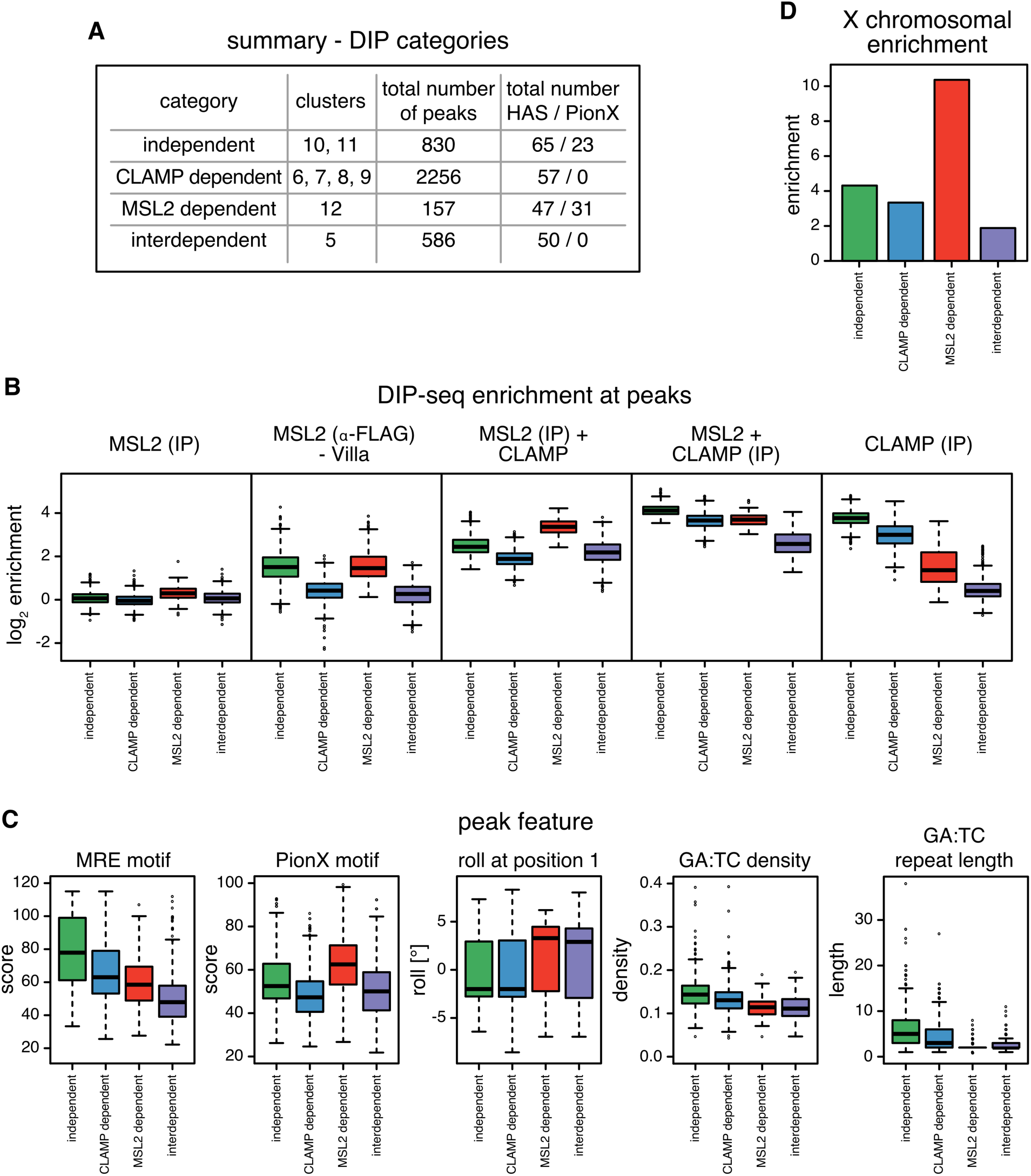
Cooperation between CLAMP and MSL2 in genome-wide DNA binding *in vitro*. **(A)** Summary of distinct binding properties discovered in the heat map (Figure 4). Clusters were assigned into four categories: independent, CLAMP-dependent, MSL2-dependent and interdependent. **(B)** Boxplot of mean log_2_ enrichment (n = 3) of *in vitro* DIP-seq at peaks grouped by the four categories described in (A) for MSL2 (IP *α*-MSL2), MSL2 and CLAMP (IP*α*-MSL2); CLAMP (IP *α*-CLAMP); MSL2 and CLAMP (IP *α*-CLAMP). **(C)** Boxplot of peak features grouped by the four categories described in (A). Panels from left to right show: score for the best matching MRE motif; score for the best matching PionX motif; the roll at position 1 of the best matching PionX motif; the density of GA:TC dinucleotides and the length of GA:TC repeats. **(D)** Bar chart of X chromosomal enrichment of peaks grouped by the four categories described in (A).

*De novo* discovery of most-enriched sequence motifs within each cluster yielded variations of the GA-rich MRE motif of variable length and regularity (Appendix S3). We attempted to stratify the binding sites further by extracting additional sequence features within each binding site (Fig. 5C, Appendix S3, S4B): (1) the score of the best-matching MRE motif; (2) the score of the best matching PionX motif; (3) the DNA roll at position +1 of the best-matching PionX motif (high roll at position ‘+1’ is a defining feature of the PionX signature (Villa et al., 2016); for simplicity we assigned the inter-base structural feature roll to the lead base pair); (4) density of GA:TC-dinucleotides; (5) length of the longest (GA:TC)_x_-repeat (as *x* repeats).

Interestingly, the ‘independent’ sites have the highest MRE scores, the highest GA:TC-dinucleotide density and the longest GA:TC-repeats (Fig. 5C, Appendix S4B). This is in good agreement with previous findings, as the first motif discovered in MSL2 *in vitro* binding sites is a long, low-complexity GA:TC-repeat, which was suggested to be bound through the MSL2 C-terminus (Villa et al., 2016), and CLAMP *in vitro* binding intensities correlate with GA:TC-repeat length (Kuzu et al., 2016). The ‘CLAMP-dependent’ sites tend to have higher MRE scores, higher GA:TC-dinucleotide density, longer GA:TC-repeats, but low PionX scores as well as low roll at positon +1. By contrast, the ‘MSL2-dependent’ sites tend to have lower MRE scores, lower GA:TC-dinucleotide density and shorter GA:TC-repeats, but the highest PionX scores including the ‘high roll at positon +1’ (Villa et al., 2016). Interestingly, ‘interdependent’ sites lack the features that characterize CLAMP or MSL2 binding sites *in vitro* (high PionX and MRE scores, high GA:TC-dinucleotide density and long GA:TC-repeats length), with the exception of ‘high roll at position +1’. Apparently, neither MSL2 nor CLAMP alone bind well to these sites, but together they mount an interaction surface able to recognizes a degenerate GA-rich MRE consensus motif (Appendix S3). We speculate that the CXC domain of MSL2 reads out the DNA shape at the 5’ end of these binding sites.

The cooperativity between MSL2 and CLAMP expands each other’s binding repertoire *in vitro*. Together both proteins are capable to identify nearly all physiological functional MRE sequencing (HAS) *in vitro*. They also bind to many autosomal sites, which may be occluded by nucleosomes *in vivo*. Accordingly and in agreement with earlier findings (Villa et al., 2016), the enrichment of X chromosomal sequences *in vitro* is almost entirely due to the action of MSL2. MSL2-dependent sites which harbor the PionX signature are 10.4-fold enriched on the X chromosome, recapitulating the 9.8-fold enrichment of PionX sequences (Fig. 5D and Appendix S4C) (Villa et al., 2016).

### CLAMP and MSL2 form a stable complex

The results of the DIP experiments suggest cooperativity between CLAMP and MSL2. Such synergism may be due to physical interaction of the two proteins. Previously, CLAMP has been found associated with the DCC after crosslinking *in vivo* (Wang et al., 2013).

As a direct test for protein interactions we employed a Yeast Two-Hybrid assay (Y2H). MSL2 (and deletion derivatives) were expressed in fusion with a Gal4 DNA binding domain (DBD) along with CLAMP (and deletion derivatives) fused to the Gal4 activation domain (AD). If co-expressed in yeast, the association of the two proteins reconstitutes the function of the transcription factor GAL4 that activates the *his3* gene in the yeast strain pJ69-4A that is auxotroph for histidine (Feilotter et al., 1994). Yeasts in which DBD and AD fusion proteins interact can grow on plates lacking histidine, leucine and tryptophan. The assay revealed a robust and reproducible interaction of MSL2-DBD and AD-CLAMP (Appendix S5A, S6A) even in stringent conditions posed by absence of adenine and presence of 3-amino-1,2,4-triazole, a competitive inhibitor of the enzyme encoded by *his3*. MSL2-DBD or AD-CLAMP alone did not support growth in the presence of the unfused AD or DBD, respectively (Appendix S5A). Using appropriate deletion constructs we found that the first 153 amino acids of CLAMP are sufficient to interact with DBD-MSL2 but further deletion to the first 123 amino acids abolished the interaction (Appendix S5A). This assay locates the MSL2 interaction site to an N-terminal fragment of CLAMP, which harbors the first ZnF domain. The remaining 6 C-terminal ZnF domains are involved in binding to GA-dinucleotide repeats (Kaye et al., 2018; Kuzu et al., 2016; Soruco et al., 2013). Using various C-terminal deletion constructs of MSL2 showed that the ∽200 C-terminal amino acids downstream of the CXC domain are required for interaction with CLAMP (Appendix S5A). As a control for proper MSL2 folding we confirmed that MSL2^1-573^-DBD was still able to interact with MSL1 via the N-terminal RING domain in our assay (Appendix S6B) as reported previously (Hallacli et al., 2012).

To probe whether this interaction was direct we tested the recombinant purified proteins used in DIP in the absence of DNA. The two proteins were probed at equimolar concentration (100 nM) for interaction by co-immunoprecipitation (co-IP) with *α*-MSL2 and *α*-CLAMP antibodies. Both proteins were quantitatively (∽90%) immunoprecipitated with the *α*-MSL2 antibody (Figs. 6A, B). While with the *α*-CLAMP antibody only ∽50% of each protein was immunoprecipitated, presumably because the antibody amount was limiting. Of note, the IP of CLAMP was more efficient in the presence of MSL2 (Figs. EV3A and B), perhaps due to conformational stabilization.

**Figure 6.**
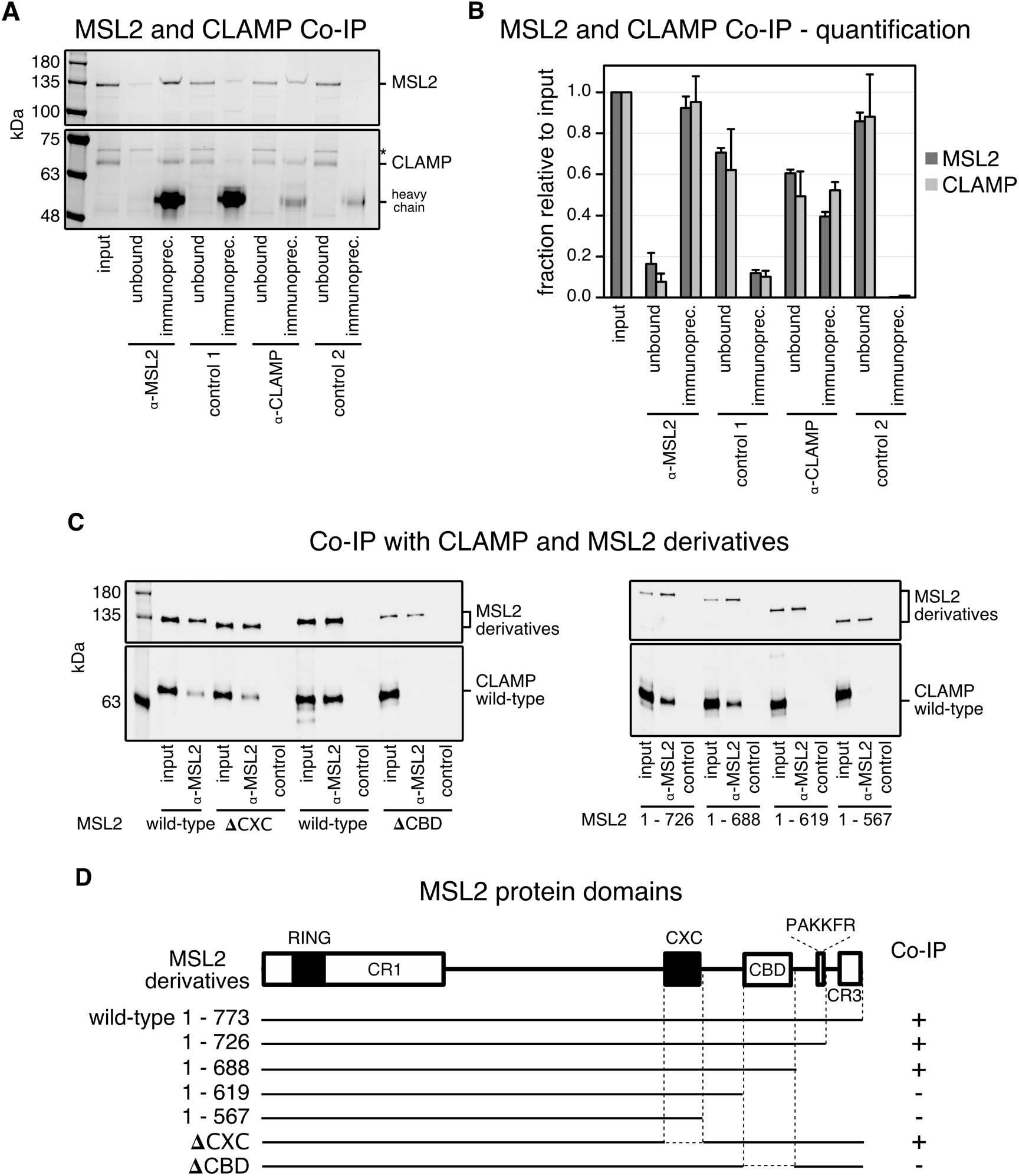
A conserved C-terminal region in MSL2 is responsible for CLAMP binding. **(A)** SDS-PAGE analysis with Coomassie staining of co-IP fractions. Purified wildtype recombinant MSL2-FLAG and CLAMP-FLAG were immunoprecipitated with *α*-MSL2 serum and the corresponding pre-immune serum as control (control 1), and with affinity-purified *α*-CLAMP antibody mixed into an irrelevant rabbit serum and with the irrelevant rabbit serum only as control 2. The corresponding unbound fractions are loaded next to each IP. A contaminant present in the MSL2 preparation is labeled with asterisk. Molecular weight markers are shown to the left. **(B)** Bar chart of the quantification from co-IP experiments as in (A), combining data from three independent MSL2-FLAG and CLAMP-FLAG purifications. The amount of each protein in the unbound fractions and IP’s were quantified relative to the input. Error bars represent the standard deviation (n = 3). **(C)** Quantitative Western blot analysis using *α*-FLAG antibody of co-IP experiments with extracts from Sf21 cells expressing wild-type CLAMP-FLAG and various MSL2-FLAG C-terminal deletion mutants. Co-IP was performed with *α*-MSL2 serum and the corresponding pre-immune serum as control [control 1 in (A)]. IP fractions were loaded next to each corresponding input. **(D)** Summary of MSL2 and CLAMP interaction from co-IP experiments presented in (C). Interactions with a CLAMP / MSL2 ratio in *α*-FLAG Western blot analysis between 0.3 to 1.7 are depicted by (+) and no detectable interaction by (-). The MSL2 domain architecture is drawn to scale. White rectangles represent the conserved regions: CR1, CBD [CR2] and CR3 (Figs. Appendix S5B and Appendix S8). Black rectangles represent the RING and CXC domains.

To map the interaction surfaces within MSL2 and CLAMP, we co-expressed deletion mutants of both recombinant FLAG-tagged proteins in Sf21 cells and immunoprecipitated them from total cell extracts. To map the CLAMP-interaction site in MSL2, we used a series of C-terminal deletions systematically lacking conserved regions identified within MSL2. In short, alignment of MSL2 sequences from 12 *Drosophila* species revealed five conserved regions (CR) (Appendix S5B, Appendix S8): the highly conserved RING and CXC domains; CR2 consisting of 66 amino acids with 50%-90% conservation; a conserved PAKKFR motif, as part of a stretch of 25 residues enriched in basic amino acids; and CR3, which corresponds to the 28 C-terminal amino acids of the *D. melanogaster* MSL2 (Appendix S5B). The 20 amino acid proline-rich region within MSL2’s C-terminus separates the CR2 from the PAKKFR motif.

MSL2 derivatives harboring CR2 (MSL2^1-726^ and MSL2^1-688^) showed quantitative binding of CLAMP, whereas fragments lacking CR2 (MSL2^1-619^ and MSL2^1-567^) did not bind CLAMP (Figs. 6C, D). CLAMP binding was unaffected by deletion of the CXC domain, but internal deletion of CR2 abolished all CLAMP binding. We therefore refer to the conserved region between amino acids 620-685 as ‘CLAMP-binding domain’ (CBD) of MSL2.

To interrogate the MSL2 interaction surface in CLAMP we used conditions where CLAMP was in excess over MSL2 to favor the interaction with MSL2 in *α*-MSL2 co-IP experiments. While full-length CLAMP reproducibly bound to MSL2, none of eight CLAMP deletion mutants, including the N-terminal CLAMP^1-153^ fragment, which interacted with MSL2 in the Y2H assay (Appendix S5), bound to MSL2 above a background, even at protein concentrations in the extracts approaching 100 nM (Figs. EV3C-F).

In summary, our data document a stable interaction between MSL2 and CLAMP and we map the interaction domains to the N-terminus of CLAMP including the first zinc finger domain and the CBD of MSL2 just downstream of the CXC domain.

### CLAMP and MSL2 cooperate to keep HAS nucleosome-free

The genome-wide DIP revealed the potential for extensive cooperation between MSL2 and CLAMP to bind shared binding sites, but the physiological X/autosome discrimination of MSL2 was not improved in free DNA. Conceivably, exclusive X chromosome binding requires a chromatin environment.

To survey the contribution of either factor to HAS accessibility in chromatin, we performed ATAC-seq after RNAi against *clamp* or *msl2* in male S2 and female Kc cells (Figs. 7A and EV4A). Depletion of CLAMP in S2 and Kc cells by RNAi caused only few significant changes in accessibility genome-wide (258 of 8913 sites and 102 of 9767 sites in S2 and Kc cells, respectively; Fig. 7B and EV4B). This indicates that most sites bound by CLAMP are kept accessible by other factors. Most sites affected by CLAMP depletion became less accessible, showing that CLAMP contributes to keeping particular loci open. These sites, that depend on CLAMP to be accessible, include especially many HAS (61 HAS and 6 HAS-PionX) in S2 cells. Focusing on the 309 HAS showed that HAS are commonly accessible in male S2, but not in female Kc cells (Fig. EV4B and C) suggesting that the DCC is required for HAS accessibility. Indeed, depletion of MSL2 in S2 cells caused only few significant changes in accessibility genome-wide (61 of 8913 sites), but these sites were nearly exclusively HAS [47 HAS and 7 HAS-PionX] (Fig. 7B).

**Figure 7.**
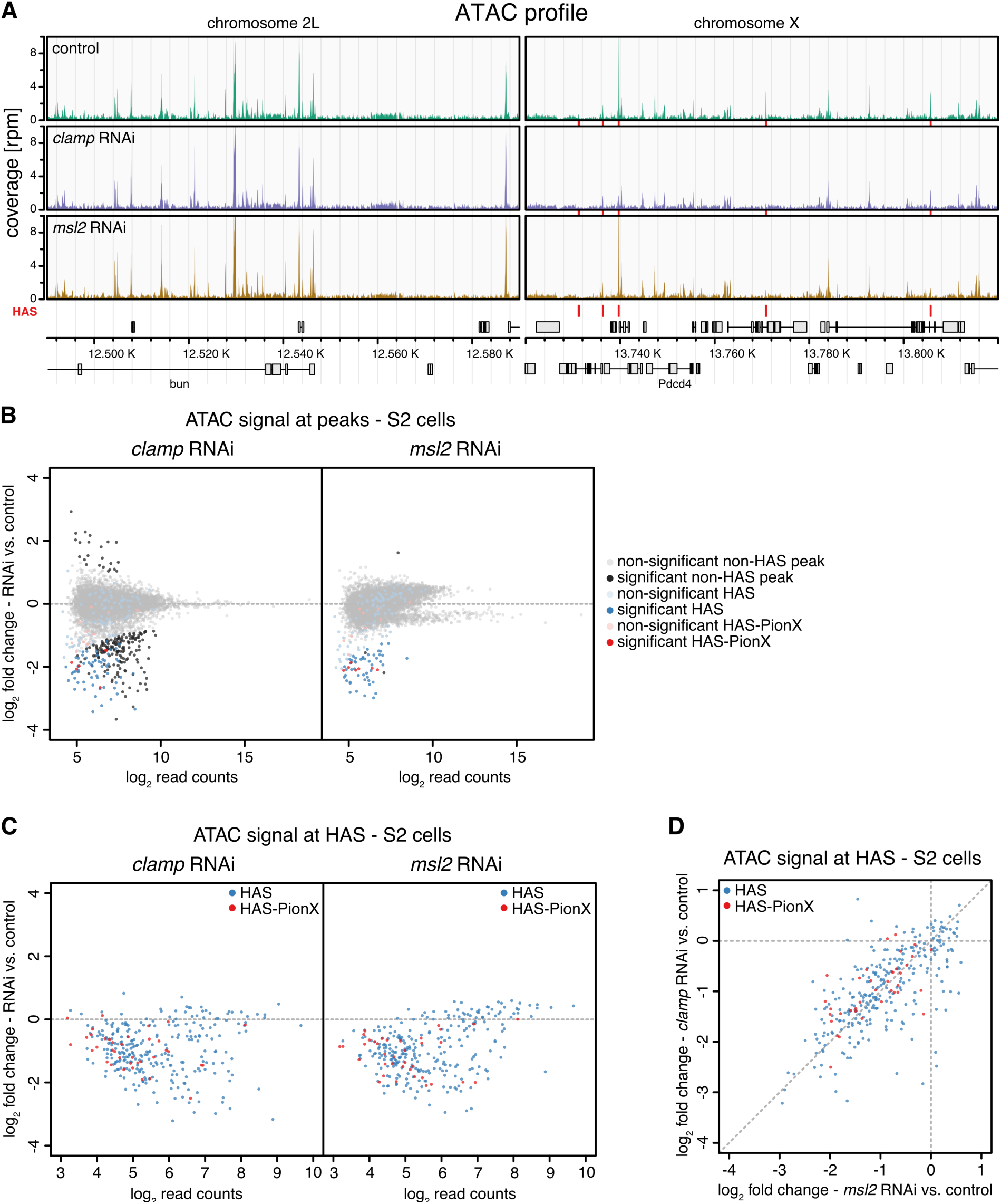
MSL2 and CLAMP synergize to keep HAS accessible in male cells. **(A)** Genome browser profile of ATAC-seq profile showing mean coverages (n = 3) along representative 100 kb windows on chromosome 2L and X. The panels show control S2 cells and cells after *clamp* RNAi or *msl2* RNAi as indicated. Red bars above the gene models and between the panels mark positions of HAS. **(B)** Scatter plot of mean log_2_ fold-change (n = 3) of ATAC-seq signal in S2 cells upon *clamp* RNAi (left panel) and *msl2* RNAi versus controls (right panel) against mean log_2_ read count in control at robust ATAC peaks (n = 8913). HAS non-overlapping with PionX sites (HAS) are marked in blue and HAS overlapping with PionX sites (HAS-PionX) are marked red (the remaining sites are displayed in grey). Sites with statistically significant different ATAC signal between RNAi and control conditions (|lfc| > 0.5 and fdr < 0.1) are marked in darker color. For *clamp* RNAi, 258 ATAC peaks are statistically significant different between conditions, including 61 HAS and 6 HAS-PionX. For *msl2* RNAi, 61 ATAC peaks are statistically significant different between conditions, including 49 HAS and 7 HAS-PionX. **(C)** Scatter plot of mean log_2_ fold-change (n = 3) of ATAC-seq signal in S2 cells upon *clamp* RNAi (left panel) and *msl2* RNAi versus controls (right panel) against mean log_2_ read count in control at HAS (n = 309). HAS non-overlapping with PionX sites (HAS, n = 272) are marked in blue and overlapping with PionX sites (HAS-PionX, n = 37) are marked red. **(D)** Scatter plot of mean log_2_ fold-change (n = 3) of ATAC-seq signal in S2 cells upon *clamp* RNAi (left panel) against *msl2* RNAi versus controls (right panel) at HAS (n = 309), as shown in (C).

Remarkably, most HAS are inaccessible in the absence of either CLAMP or MSL2 in S2 cells (Fig. 7C). Apparently, CLAMP and MSL2 contribute equally to HAS accessibility (Fig. 7D). The minority of HAS that remained accessible in the absence of MSL2 or CLAMP are also accessible in Kc cells (Figs. 7C, EV4B, C), indicating that these HAS are kept open by other factors.

Broadening the view, we found that CLAMP promotes access to other loci in the genomes of both cell lines (Fig. 7B and EV4B). According to the 9-state chromatin model (Kharchenko et al., 2011) (Fig. EV4D) many of these sites bear promoter or enhancer signatures. Unexpectedly, we found that sites marked by H3K27me3, the polycomb signature, are most enriched within the sites affected by CLAMP depletion.

These experiments identify a second layer of DNA binding ccoperativity between CLAMP and MSL2. Faced with purified genomic DNA both proteins cooperate to bind to essentially all sites with appropriate DNA sequence features, regardless of chromosomal origin. By contrast, in a physiological chromatin environment extensive cooperativity between both factors is restricted to X chromosomal HAS.

## Discussion

The process of dosage compensation in *Drosophila* provides an excellent opportunity to study the principles of sequence-selective DNA binding. Male flies only survive if the regulatory DCC exclusively binds to the X chromosome. Earlier work suggested that this exclusivity is at least partly due to the functional cooperation between the DNA-binding subunit of the DCC, MSL2 (Fauth et al., 2010; Straub et al., 2013; Villa et al., 2016; Zheng et al., 2014), and the zinc finger protein CLAMP (Kuzu et al., 2016; Larschan et al., 2012; Soruco et al., 2013). Curiously, both factors share the intrinsic property to bind to GA-rich MRE sequences that are hallmarks of the X chromosomal high affinity sites (HAS) of the DCC. We recently found that the CXC domain of MSL2 can read the DNA signature of a prominent subset of MREs with notable 5’ extension, termed PionX (Villa et al., 2016). Using chromatin immunoprecipitation with high-throughput sequencing (ChIP-seq) we now found that essentially all MSL2 interactions with HAS requires the presence of CLAMP, suggesting a tight cooperativity between both factors.

Cooperative DNA binding of transcription factors (TFs) for refined and stable recognition of complex DNA elements is a widely used principle of gene regulation (Morgunova and Taipale, 2017). Such cooperativity may involve a direct contact between two factors. Their simultaneous contact with DNA and dimerization partner reduces the off-rate of each individual binder significantly (Jolma et al., 2015). We indeed found a physical interaction between CLAMP and MSL2 and mapped the CLAMP-binding domain on MSL2 just C-terminal to the DNA-binding CXC domain.

The dimerization of TFs may happen in solution or be promoted by a target DNA, which defines spacing and orientation of the proteins binding to adjacent sequences. Although we demonstrated stable, soluble complex formation between CLAMP and MSL2, we consider it unlikely that such diffusible complexes are abundant in cells. Interactions have so far only been observed in ChIP followed by mass spectrometry with prior crosslinking (Wang et al., 2013). Probing the dynamics of MSL2 by FRAP (fluorescence recovery after photobleaching) we found earlier (and recently confirmed) that MSL2 binds the X chromosome very tightly with no evidence for a freely diffusible component (Straub et al., 2005). Further, MSL2 is only expressed during S phase, when the newly synthetized X chromosome needs to be dosage compensated (Lim and Kelley, 2012). We thus favor the idea that the proteins encounter each other on DNA and then ‘lock in’ to stabilize each other on the HAS binding site.

We speculate that this interaction may align the DNA-binding CXC domain of MSL2 with the GA-repeat binding zinc fingers of CLAMP to form a long, contiguous DNA interaction surface suited to read out the long (∽20 bp) MRE/PionX sequences. In such an arrangement, the CXC domain may recognize the prominent DNA shape feature akin to the PionX signature at the 5’ end of a binding site, whereas CLAMP may use its zinc fingers to interact with multiple GA-repeats in the 3’ part.

Our DIP analysis revealed several clusters of sequence elements that differ in MSL2 and CLAMP binding affinity and cooperativity. The different binding scenarios suggest that the cooperativity depends on the precise properties of each individual DNA sequence, such as length, composition and shape. The long GA-repeat that characterizes MREs suggests a flexible interaction of GA-recognition surfaces of either MSL2 or CLAMP to move sideways to accommodate binding of the partner. It is possible that CLAMP does not use all six GA-repeat binding zinc fingers simultaneously at shorter sites. Previously it was shown that the multiple-zinc finger protein CTCF makes flexible use of its zinc fingers to bind a diverse range of sequences (Nakahashi et al., 2013).

Although the results suggest a certain flexibility of factor interaction with DNA, some simple trends can be seen. CLAMP binding correlates with GA:CT-dinucleotide density and -repeat length and MSL2 retrieves the PionX signature, in agreement with previous observations (Kuzu et al., 2016; Villa et al., 2016). Sites that are bound in an ‘MSL2-dependent’ mode contain PionX sequences with high roll at position +1 and contribute most to X chromosome specificity, in line with our earlier conclusion (Villa et al., 2016). Furthermore, both proteins can independently bind to sequences with high MRE score and long GA-repeats, in agreement with the previous finding that also MSL2 is capable to retrieve long GA-repeats most likely through interaction via its proline-rich C-terminus (Villa et al., 2016). Conversely, sites with low MRE scores (but high roll at position +1 as characteristic for the PionX signature) and short GA-repeats can only be bound if both factors cooperate (interdependence).

TFs may cooperate for DNA binding even without direct interaction if their binding to DNA competes with nucleosome formation. In this scenario, binding of each individual TF hinders nucleosome formation and increases the likelihood that the second TF finds its close-by binding site accessible (Adams and Workman, 1995; Mirny, 2010; Polach and Widom, 1996). Our ATAC-seq study suggests that this is an important aspect of MSL2/CLAMP function. TF cooperativity may be particularly important if the concentrations of a partner is too low to effectively compete with nucleosome formation by itself (Li et al., 2011). The concentrations of MSL2 are suggested to be relatively low and tightly controlled because excess of MSL2 over X chromosomal binding sites will lead to binding of autosomal sites causing male lethality (Villa et al., 2012). Conceivably, the abundant CLAMP was coopted during the evolution of the MSL2-MRE/PionX interaction to increase the affinity of MSL2 to relevant binding sites. CLAMP binds longer GA-repeat sequences (Kuzu et al., 2016) and because its binding has been reported to lead to regions of enhanced accessibility of chromatin in its neighborhood (Urban et al., 2017a), we hypothesized that CLAMP may fulfill a ‘chromatin opener’ function for MSL2. This ‘division of labor’ model between an abundant ‘chromatin opener’ and a TF that profits from the accessible region was developed following the observation that binding of the GAGA factor (GAF) to GAGAG sequences in promoters, enhancers and polycomb response elements, leads to chromatin opening and facilitated the binding of other proteins in the neighborhood (Leibovitch et al., 2002; Lu et al., 1993; Strutt et al., 1997; Tsukiyama et al., 1994). Our data are not consistent with such a hierarchical model. Rather surprisingly, they reveal that both factors contribute equally to keep shared binding sites nucleosome-free. While CLAMP is required to stabilize the interaction of MSL2 at HAS, the reverse is also true: stable interaction of CLAMP at HAS critically depends on MSL2. Our findings are reminiscent of the recently proposed cooperativity between the pioneer factor Zelda and the morphogen Bicoid during *Drosophila* preblastoderm development (Hannon et al., 2017).

Given the many other CLAMP binding sites in the genome, we assume that CLAMP cooperates with other proteins elsewhere in the genome. Interestingly, we found that depletion of CLAMP leads to diminished accessibility at chromatin with hallmarks of polycomb repression (Fig. EV4D) may pointing to a hitherto unappreciated cooperation with the polycomb machinery. The accessibility of most other CLAMP binding sites is independent of CLAMP, suggesting that other DNA-binding proteins promote accessibility of these binding sites.

Our DIP analysis clearly documents the potential for cooperative DNA binding genome-wide, but this alone is not sufficient to discriminate the physiological HAS on the X chromosome from similar sequences that are not *in vivo* targets of the DCC. *In vivo,* the synergism between MSL2 and CLAMP only manifests in chromatin at a relatively small number of X chromosomal HAS. Our study suggests that competing chromatin assembly may pose stringent demands on the cooperative action of MSL2 and CLAMP. Additional cooperativity may manifest at the level of MSL proteins and HAS DNA. *In vivo*, assembly of MSL2 with other MSL proteins and *roX* RNA involves dimerization via the MSL1 subunit. Such dimerization will bring two DNA-binding domains of MSL2 into proximity, increasing the potential for cooperative effects. The potential of DNA recognition by combinations of MSL2 and CLAMP DNA-binding domains is matched by the complexity of their DNA targets as many High Affinity Sites contain several MRE sequences (Alekseyenko et al., 2008; Straub et al., 2008). Our systematic analysis of intrinsic DNA-binding properties of two key factors involved in X chromosome recognition and of their *in vivo* chromatin interactions sheds light on the sophistication of combinatorial DNA recognition that evolved to prevent male lethality.

## Materials and Methods

### Cell culture and RNAi

S2 (DGRC stock # 181), Kc167 (DGRC stock # 1) and L2-4 (S2 subclone, provided by P. Heun) cells were cultured in Schneider’s *Drosophila* Medium (Thermo Fisher), supplemented with 10% heat-inactivated Fetal Bovine Serum (Sigma-Aldrich), 100 units/mL penicillin and 0.1 mg/mL streptomycin (Sigma-Aldrich) at 26°C. RNAi against target genes in S2, L2-4 and Kc cells was performed as previously described (Villa et al., 2016). In brief, double-stranded RNA fragments (dsRNA) were generated with MEGAscript kit (Thermo Fisher) from PCR products obtained using the following forward and reverse primers (separated by comma): *clamp* RNAi #1: TAATACGACTCACTATAGGGACGTCCAAACCCTTCAGTTGT, TAATACGACTCACTATAGGGATTGAGTGCAAAACGATCAGC;

*clamp* RNAi #2 (DRSC29935): TTAATACGACTCACTATAGGGAGAGAAGACCTTACCAAAAACAT, TTAATACGACTCACTATAGGGAGAGCTTATGTTGGATATGGTGT; *trl* RNAi: TTAATACGACTCACTATAGGGAGAATGTCGCTGCCA, TTAATACGACTCAC-TATAGGGAGATTGCCTGGA; gst RNAi: TTAATACGACTCACTATAGGGAGAATGTCCCCTATACTAGGTTA, TTAATACGACTCACTATAGGGAGAACGCATCCAGGCACATTG; The sequence of *Schistosoma japonicum* GST was amplified from pGEX-6P-1 (GE Healthcare). *gfp* RNAi: TTAATACGACTCACTATAGGGTGCTCAGGTAGTGGTTGTCG, TTAATACGACTCACTATAGGGCCTGAAGTTCATCTGCACCA; *msl2* RNAi: TTAATACGACTCACTATAGGGAGAATGGCCCAGACGGCATAC, TTAATAC-GACTCACTATAGGGAGACAGCGATGTGGGCATGTC.

Cells were washed with serum-free medium and 10 µg dsRNA per 10^6^ cells at a concentration of 10 µg/mL in serum-free medium (10^6^ cells in 6-well plate for ATAC-seq) or 4.2 µg dsRNA per 10^6^ cells at 6.3 µg/mL serum free medium (12 * 10^6^ cells in 75 cm^2^ flask for ChIP-seq) was added, incubated for 10 min at room temperature (RT) with slight agitation and further 50 min at 26°C. Two volumes of complete growth medium were added and cells were incubated for 5 days at 26°C. Then, 1x of initial volume (6-well plate) or 0.75x of initial volume (75 cm^2^ flask) of growth medium was added and cells were incubated for further 2 days at 26°C.

Sf21 cells (Thermo Fischer) were cultured in SF900 II SFM (Thermo Fisher), supplemented with 10% heat-inactivated Fetal Bovine Serum (Sigma-Aldrich), 0.1 mg/mL gentamicin (Sigma-Aldrich) at 26°C.

### Immunofluorescence microscopy

For immunofluorescence microscopy (IFM), 0.5 * 10^6^ L2-4 cells after RNAi treatment in 200 – 500 µL growth medium were seeded into each well of a 3-well (14 mm) object slide (Thermo Fisher) and incubated for 2 h at 26°C. Immunofluorescence staining was performed as described in (Schunter et al., 2017) with slight modifications. Briefly, cells were washed in PBS, fixed for 7.5 min with 2% (v/v) paraformaldehyde (PFA) in PBS on ice and permeabilized for 7.5 min with 1% (v/v) PFA in PBS with 0.25% (v/v) Triton-X 100 on ice. Cells were washed twice in PBS and blocked for 1 h with 200 µL Blocking Buffer (3% (w/v) BSA, 5% (v/v) heat-inactivated fetal bovine serum (Sigma-Aldrich) in PBS) at RT in a humid chamber. Cells were incubated for 1 h with primary antibody diluted in 30 µL Blocking Buffer with 0.1% (v/v) Triton-X 100 at RT in a humid chamber. Cells were washed twice for 5 min in 200 µL PBS and incubated for 1 h with suitable secondary antibody diluted in 30 µL Blocking Buffer with 0.1% (v/v) Triton-X 100 at RT in a humid chamber. Cells were washed twice for 5 min in 200 µL PBS, stained for 2 min with 0.125 µg/mL DAPI in 200 µL PBS at RT and washed twice for 5 min in 200 µL PBS. Cells were mounted with 8 µL Vectashield (Vector Laboratories) and a coverslip was sealed to object slide with nail polish. Images were acquired with Axiovert 200 epifluorescence microscope (Zeiss) equipped with AxioCamMR CCD Camera (Zeiss).

### Protein Purification

Recombinant MSL2-FLAG was expressed in Sf21 cells and purified by FLAG-tag affinity chromatography, as previously described in (Fauth et al., 2010). In brief, Sf21 cells at 10^6^ cells/mL (250 * 10^6^ cells) were infected 1:1000 (v/v) with baculovirus, expressing MSL2-FLAG. After 72 h, cells were harvested and washed once in PBS, frozen in liquid nitrogen and stored at −80°C. To lyse, cells were rapidly thawed, resuspended in 1 mL Lysis Buffer per 10 mL of culture (50 mM HEPES pH 7.6, 300 mM KCl, 1 mM MgCl_2_, 5% (v/v) glycerol, 0.05% (v/v) IGEPAL CA-360, 50 µM ZnCl_2_) supplemented with 0.5 mM TCEP and cOmplete EDTA-free Protease Inhibitor Cocktail (Sigma-Aldrich) (PI). The suspension was passed thrice through Microfluidizer LM10 (Microfluidics). Cell extract was adjusted with Lysis Buffer containing 0.5 mM TCEP and PI to 2 mL per 10 mL of culture and incubated with end-over-end rotation for 1 h at 4°C. Cell debris were spun down at 4°C for 30 min at 50,000 g. The resulting supernatant was used for FLAG-tag affinity purification with 4 µL of a 50% slurry of FLAG-M2 beads (Sigma-Aldrich) per 1 mL of culture. Beads were first washed thrice in 20 bed volumes of Lysis Buffer, subsequently supernatant was added and incubated with end-over-end rotation for 3 h at 4°C. Beads were pelleted (4°C, 5 min, 500 g) and supernatant was removed. Beads were washed once with 20 bed volumes each of Lysis Buffer, Wash Buffer (Lysis Buffer with 1000 mM KCl), Lysis Buffer again, and finally twice with 20 bed volumes Elution Buffer (Lysis Buffer with 100 mM KCl). For protein elution, beads were incubated with 0.2 bed volumes of Elution Buffer containing 5 mg/mL FLAG peptide for 10 min at 4°C and subsequently 1.8 bed volumes of Elution Buffer with 0.5 mM TCEP and PI were added and incubated with end-over-end rotation for 30 min at 4°C. The elution step was repeated once. Elution fractions were combined, remaining beads were removed by passing through Corning Costar Spin-X centrifuge tube filters (Sigma-Aldrich) and the eluate was concentrated with 10 MWCO Amicon Ultra 0.5 mL (Merck). Protein concentration was determined using BSA standards on SDS-PAGE with Coomassie brilliant blue G250 staining. To store at −80°C until use, concentrated protein was aliquoted and flash-frozen in liquid nitrogen.

For purification of recombinant CLAMP by FLAG-tag affinity chromatography, the coding sequence of CLAMP (CG1832-RA) was fused to a C-terminal coding sequence of FLAG affinity tag and cloned into pFastBac1, using the following forward and reverse primer: AAGGATCCATGGAAGACCTTACCAAAAAC, CCTTTCTCGAGTTACTT-GTCATCGTCGTCCTTGTAGTCTTCCCCGTCTGTATGCATCCG.

CLAMP-FLAG was expressed in Sf21 cells and purified by FLAG-tag affinity chromatography, like MSL2-FLAG, but with the following modifications. Lysed cells were resuspended in 1 mL Buffer C per 10 mL of culture (50 mM HEPES pH 7.6, 1 M KCl, 1 mM MgCl_2_, 5% (v/v) Glycerol, 0.05% (v/v) IGEPAL CA-360, 50 µM ZnCl_2_, 375 mM L-Arginine (according to (Leibly et al., 2012)) supplemented with 0.5 mM TCEP and cOmplete EDTA-free Protease Inhibitor Cocktail (Sigma-Aldrich) (PI) and passed thrice through Microfluidizer LM10 (Microfluidics). Subsequently, the extract was adjusted with Buffer C containing PI to 2 mL per 10 mL of culture and supplemented with 0.1% (v/v) polyethylenimine by adding 2% (v/v) polyethylenimine (neutralized with HCl to pH 7.0) drop-by-drop while string in an ice bath, according to (Patel et al., 2016). Cell debris were spun down at 4°C for 30 min at 50,000 g. To the resulting supernatant 20 U Benzonase was added per 10 mL of culture and incubated with end-over-end rotation for 1 h at 4°C. The resulting extract was used for FLAG-tag affinity purification with 4 µL of a 50% slurry of FLAG-M2 beads (Sigma-Aldrich) per 1 mL of culture. Beads were washed thrice in 20 bed volumes of Buffer C, subsequently supernatant was added and incubated with end-over-end rotation for 3 h at 4°C. Beads were pelleted at 4°C for 5 min at 500 g and supernatant was removed. Beads were washed 5 times with 20 bed volumes of Buffer C. For protein elution, beads were incubated with 0.2 bed volumes of Buffer C containing 5 mg/mL FLAG peptide for 10 min at 4°C and subsequently 1.8 bed volumes of Buffer C containing 0.5 mM TCEP and PI were added and incubated with end-over-end rotation for 30 min at 4°C. The elution step was repeated once. The combined elution fractions were processed on flash-frozen in liquid nitrogen as for MSL2-FLAG.

### Genomic DNA preparation

For genomic DNA (gDNA) extraction, 5 * 10^7^ S2 cells were harvested, washed in PBS and gDNA was extracted with NucleoSpin Tissue kit (Macherey-Nagel). Remaining RNA contaminants in eluted gDNA were digested with 0.1 mg/mL RNaseA (Sigma-Aldrich) for 1 h at 37°C. Subsequently, gDNA was sonicated with Covaris AFA S220 using microTUBEs at 175 W peak incident power, 10% duty factor and 200 cycles per burst for 430 s at 5°C to generate ∽150-200 bp fragments, as described in (Villa et al., 2016). Sheared gDNA was purified with MinElute kit (QIAGEN), concentration was determined using Qubit (Thermo Fisher) and fragment size was verified using 2100 Bioanalyzer (Agilent).

### Antibodies

For immunoblotting, affinity-purified polyclonal *α*-H3 C-term antibody (Abcam, ab1791), affinity-purified polyclonal *α*-CLAMP antibody (Novus, 49880002), polyclonal rabbit *α*-MSL2^Gilfillan^ serum [‘*α*-MSL2^Gilfillan^’ (Gilfillan et al., 2006)], polyclonal rabbit *α*-MSL2 serum [generated by Pineda Antikörper-Service here termed ‘*α*-MSL2^Pineda^’ against the MSL2-fragment (amino acids 296-608) similar to the one used for *α*-MSL2^Gilfillan^, polyclonal rabbit *α*-GAF serum [‘*α*-GAF’ (Strutt et al., 1997)] and affinity-purified monoclonal *α*-FLAG M2 antibody (Sigma-Aldrich, F3165) were used. For immunostaining, *α*-MSL2^Gilfillan^ and culture supernatant containing monoclonal *α*-MSL3 (Straub et al., 2008) were used. For ChIP, *α*-MSL2^Gilfillan^, *α*- GAF and *α*-CLAMP antibodies were used. For DIP, supernatant containing monoclonal *α*-MSL2 [generated by E. Kremmer (Helmholtz Zentrum Munich, Germany) against the same MSL2-fragment as used for *α*-MSL2^Gilfillan^] and supernatant containing monoclonal *α*-CLAMP [generated by E. Kremmer (Helmholtz Zentrum Munich, Germany) against the peptide LATTDDNKTCYI] were used. For IP, serum containing polyclonal *α*-MSL2^Pineda^ and affinity-purified polyclonal *α*-CLAMP* antibody described in (Rieder et al., 2017) (kind gift of E. Larschan) were used.

### Yeast two-hybrid assay

Yeast two-hybrid assay was carried out using yeast strain pJ69-4A (MATa trp1-901 leu2-3,112 ura3-52 his3-200 gal4Δ gal80Δ GAL2-ADE2 LYS2::GAL1-HIS3 met2::GAL7-lacZ), with plasmids and protocols from Clontech. For growth assays, plasmids were transformed into yeast strain pJ69-4A by the lithium acetate method, as described by the manufacturer with some modifications. All cells were grown at 30°C in an orbital shaker at 250-300 rpm. Yeast colonies were transferred into a 15 mL culture tube with 6-7 mL of YPDA medium and grown for one day. The culture was 10-fold diluted in a 0.5 L culture flask with YPDA medium and cultivated for 3 h. Aliquots of 1.5 mL cell suspension were pelleted by centrifugation at 4000 g for10 s, and the supernatant was removed. Pelleted cells were resuspended in 1 mL 0.1 M LiAcO, incubated for 30 min and pelleted as above. To the pellet, 240 μL 50% (v/v) PEG 3380, 36 μL 1 M LiAcO, and 50 μL of a mixture of two plasmids (the amount of each plasmid in a mixture of 400-800 ng) was added sequentially and the cells suspended to homogeneity. The tube was incubated at 30°C for 30 min, then at 42°C for 5 min and then placed on ice for 1-2 minutes. Cells were pelleted at 4000 g for 15-20 s and resuspended in 100 μL ddH_2_O. The resuspended cells were plated on selective medium lacking Leu and Trp (“medium-2”). The plates were incubated at 30°C for 2-3 days. Afterwards, the colonies were streaked out on plates on selective medium lacking either Leu, Trp and His “medium-3”), or lacking adenine in addition to the three amino acids (“medium-4”), or lacking the three amino acids but containing 5 mM 3-amino-1,2,4-triazole (“medium-3 + 5mM 3AT”). The plates were incubated at 30°C for 3-4 days and growth assessed. Each assay was prepared as three independent biological replicates with three technical repeats. Fusion proteins were cloned into pGBT9 and pGAD424 vectors from Clontech (Appendix Table S1) and verified by sequencing.

### CLAMP-MSL2 co-immunoprecipitation

Per reaction, 100 nM recombinant MSL2-FLAG and CLAMP-FLAG in 50 µL Binding Buffer (2 mM Tris/HCl pH 7.5, 100 mM KCl, 2 mM MgCl_2_, 10% (v/v) Glycerol, 10 µM ZnCl_2_) were incubated with end-over-end rotation for 30 min at 26°C. For *α*-MSL2 immunoprecipitation, the reaction was added to 25 µL magnetic Dynabeads Protein G (Thermo Fisher) pre-coupled with antibodies. For *α*-CLAMP immunoprecipitation, the reaction was added to 5 µL beads pre-coupled with antibodies. For pre-coupling, beads were washed thrice with 1 mL Binding Buffer and incubated with end-over-end rotation for over-night at 4°C with 1 µL*α*-MSL2 antibody per 10 µL beads or with the corresponding pre-immune serum as control and with 20 µL *α*-CLAMP antibody complemented with 0.75 µL of an irrelevant pre-immune serum per 10 µL beads or with 1µL irrelevant pre-immune serum as control. Beads were washed thrice with 1 mL Binding Buffer, the co-IP reaction was added and incubated with end-over-end rotation for 15 min at RT. Beads were washed thrice in 100 µL Binding Buffer, resuspended in SDS-Sample Buffer and analyzed together with the corresponding input and unbound sample by SDS-PAGE with Coomassie brilliant blue G250 staining, quantified using a ChemiDoc Touch Imaging System (Bio-Rad). Co-IP reactions were performed with three independent CLAMP and MSL2 preparations.

### Mapping of interaction domain

Sf21 cells at 10^6^ cells/mL (20 * 10^6^ cells) were infected 1:1000 (v/v) with each baculovirus stock separately. While MSL2^ΔCXC^-FLAG and MSL2-FLAG were described previously in (Fauth et al., 2010), new constructs have been generated for CLAMP-FLAG (see above) and deletion mutants of MSL2 and CLAMP, cloned into into pFastBac1 using the corresponding primer (Appendix Table S2). After 72 h, cells were harvested and lysed in 1 mL Lysis Buffer (50 mM HEPES pH 7.6, 300 mM KCl, 1 mM MgCl_2_, 5% (v/v) glycerol, 0.05% (v/v) IGE-PAL CA-360, 50 µM ZnCl_2_) with cOmplete EDTA-free Protease Inhibitor Cocktail (Sigma-Aldrich) (PI) per 20 mL of culture. The relative amounts of expressed protein were quantified by *α*-FLAG western blot. Per co-IP reaction, MSL2-FLAG and CLAMP-FLAG (or their mutants) containing extracts were mixed in 1:1 to 1:3 ratio and if required adjusted with cell extract from uninfected cells. Cell extracts were supplemented with 12.5 U Benzonase, 1 mM DTT, 0.2 mM MgCl2 and PI, incubated with end-over-end rotation for 10 min at RT and cell debris were spun down at 4°C for 10 min at 20,000 g. For *α*-MSL2 immunoprecipitation, the reaction was added to 15 µL magnetic Dynabeads Protein G (Thermo Fisher) pre-coupled with antibodies. For pre-coupling, beads were washed thrice with 1 mL Lysis Buffer and incubated with end-over-end rotation for over-night at 4°C with 1 µL *α*-MSL2 antibody per 10 µL beads or with the corresponding pre-immune serum as control. Beads were washed thrice with 1 mL Lysis Buffer, the extract mixture was added and incubated with end-over-end rotation for 45 min at 4°C. Beads were washed thrice in 500 µL Binding Buffer, resuspended in SDS-Sample Buffer and analyzed together with the corresponding input sample by SDS-PAGE and *α*-FLAG western blot.

### ATAC-seq

ATAC-seq was performed as described earlier (Buenrostro et al., 2013; Buenrostro et al., 2015) with some modification for *Drosophila* cells. In brief, 50,000 S2 or Kc cells, also with prior RNAi, were used per reaction. Cells were washed in 100 µL PBS, resuspended in 100 µL Lysis Buffer (10 mM Tris/HCl pH 7.4, 10 mM NaCl, 3 mM MgCl_2_, 0.1% (v/v) IGE-PAL CA-630) with cOmplete EDTA-free Protease Inhibitor Cocktail (Sigma-Aldrich) and incubated for 3 min on ice. Nuclei were spun down at 4°C for 10 min at 600 g, resuspended in 50 µL 1x TD buffer with 2.5 µL TD Enzyme (Illumina) and incubated with slight agitation for 30 min at 37°C. Tagmented DNA was purified with MinElute kit (QIAGEN) and eluted in 15 µL H_2_O. To determine the cycle number for library amplification, quantitative PCR was performed in triplicates with 0.5 µL sample in 10 µL reaction containing 1x NEBNext HiFi PCR mix (NEB), 1.25 µM Ad1 and Ad2 primer each and 0.5x SYBRGreen (Thermo Fisher). Cycle number at quarter maximal intensity was used for library amplification (typically 11-13 cycles). Libraries were amplified by using 12.5 µL sample in 50 µL PCR reaction containing 1x NEBNext HiFi PCR mix (NEB), 1.25 µM Ad1 and Ad2 primer each. ATAC libraries were purified with MinElute kit (QIAGEN) and concentration determined using 2100 Bioanalyzer with High Sensitivity DNA kit (Agilent). Libraries were sequenced on HiSeq 1500 (Illumina) instrument yielding typically 25-35 million 50 bp paired-end reads per sample.

### ChIP-seq

ChIP-seq was performed as previously described (Straub et al., 2013) with slight modification. S2 cells (∽10^8^ cells) after RNAi were harvested and chilled on ice. Cells were cross-linked with 1% formaldehyde for 55 min on ice by adding 1 mL volume 10x fixing solution (50 mM HEPES pH 8.0, 100 mM NaCl, 1 mM EDTA, 0.5 mM EGTA) with 10% formaldehyde per 10 mL culture and reaction was stopped by adding 125 mM glycine and incubating for 10 min on ice. For nuclei isolation, cells were washed in PBS and resuspended in Buffer A (10 mM Tris/HCl pH 8.0, 10 mM EDTA, 0.5 mM EGTA, 0.25% (v/v) Triton-X 100) with cOmplete EDTA-free Protease Inhibitor Cocktail (Sigma-Aldrich) (PI) at 10^8^ cells/mL and incubated with end-over-end rotation for 10 min at 4°C. The cells were pelleted at 4°C for 10 min at 2300 g and resuspended in Buffer B (10 mM Tris/HCl pH 8.0, 200 mM NaCl, 1 mM EDTA, 0.5 mM EGTA, 0.01% (v/v) Triton-X 100) with PI at 10^8^ cells/mL and incubated with end-over-end rotation for 10 min at 4°C. For chromatin fragmentation, nuclei were spun down at 4°C for 10 min at 2300 g, resuspended in RIPA (10 mM Tris/HCl pH 8.0, 140 mM NaCl, 1 mM EDTA, 1% (v/v) Triton-X 100, 0.1%(v/v) SDS, 0.1% (v/v) DOC) with PI at 10^8^ cells/mL in 1 mL for shearing with Covaris AFA S220 using 12×12 tubes at 100 W peak incident power, 20% duty factor and 200 cycles per burst for 25 min at 5°C to generate 150-200 bp fragments. Protein A and Protein G (GE Healthcare) beads (mixed in a 1:1 ratio) were washed thrice with 30 bed volumes RIPA. To remove cell debris, sheared chromatin was centrifuged at 4°C for 15 min at 15,000 g and 100 µL soluble chromatin in the supernatant was pre-cleared with 3 µL (6 µL 50% slurry) Protein A and Protein G beads mix by incubating with end-over-end rotation for 1 h at 4°C. Beads were pelleted at 4°C for 5 min at 500 g and supernatant was directly used for immunoprecipitation by adding antibody to 200 µL chromatin adjusted to 500 µL with RIPA including PI and incubating with end-over-end rotation for 16 h at 4°C. To remove precipitates, chromatin was centrifuged at 4°C for 15 min at 15,000 g and 100 µL supernatant was added to 3 µL Protein A and Protein G beads mix (RIPA-equilibrated as above) by incubating with end-over-end rotation for 4 h at 4°C. Beads were spun down and washed five times with 60 bed volumes RIPA including PI by incubating with end-over-end rotation for 10 min at 4°C. For DNA recovery, beads were spun down, resuspended in 6.7 bed volumes TE Buffer (10 mM Tris/HCl pH 8.0, 1 mM EDTA), RNA was digested with 50 µg/mL RNaseA (Sigma-Aldrich) for 30 min at 37°C, and after addition of 0.5% (m/v) SDS, proteins were digested with 0.5 µg/mL Proteinase K (Sigma-Aldrich) for 16 h at 65°C with agitation. DNA was purified with 1.8x AMPure XP beads (Beckmann Coulter). Quantitative PCR was performed using SYBR Green PCR Master Mix (Thermo Fisher) and 0.5 nM forward and reverse primer each (Appendix Table S3), analyzed on LightCycler 480 (Roche). Libraries were prepared with NEBNext ChIP-Seq Library Perp Kit for Illumina (NEB) and analyzed with 2100 Bioanalyzer with DNA 1000 kit (Agilent). Libraries were sequenced on HiSeq 1500 (Illumina) instrument yielding typically 25-30 million 50 bp single-end reads per sample.

### DIP-seq

DIP-seq was performed accordingly to (Villa et al., 2016), with modifications for combined DIP of two proteins. Sheared gDNA was diluted to 4 mg/mL in Binding Buffer (2 mM Tris/HCl pH 7.5, 100 mM KCl, 2 mM MgCl_2_, 10% (v/v) Glycerol, 10 µM ZnCl_2_). Ten microliter (corresponding to 10%) was taken as input material and adjusted to 50 µL with Binding Buffer for DNA purification. Per reaction, 100 nM recombinant MSL2 and/or 50 nM recombinant CLAMP was added in 100 µL diluted gDNA and incubated with end-over-end rotation for 30 min at 26°C. For immunoprecipitation, the reaction was added to 7.5 µL (15 µL 50% slurry) Protein G beads (GE Healthcare) pre-coupled with antibodies. For pre-coupling, beads were washed thrice with 100 µL Binding Buffer and incubated with end-over-end rotation for 3-4 h at 4°C with 1 mL culture supernatant containing monoclonal antibody or culture medium as control. Beads were spun down for 1 min at 500 g, washed thrice with 100 µL Binding Buffer, the DIP reaction was added and incubated with end-over-end rotation for 15 min at RT. Beads were spun down for 1 min at 500 g, washed twice in 100 µL Binding Buffer and resuspended in 50 µL Binding Buffer. Five microliter were taken for Western blot analysis. Proteins were digested by adding Proteinase K at 0.5 mg/mL and incubated for 1 h at 56°C with agitation and DNA was purified with 1.8x AMPure XP beads (Beckmann Coulter). Libraries were prepared with MicroPlex Library Preparation Kit v2 (Diagenode) and analyzed with 2100 Bioanalyzer with DNA 1000 kit (Agilent). Libraries were sequenced on HiSeq 1500 (Illumina) instrument yielding typically 20-40 million 50 bp single-end reads per sample. DIP reactions were performed with three independent CLAMP and MSL2 preparations.

### Data analysis

Sequencing data were processed using SAMtools (Li et al., 2009) version 1.3.1, BEDtools (Quinlan and Hall, 2010) version 2.26.0, R version 3.4.2 (http://www.r-project.org) and Bioconductor (http://www.bioconductor.org) using default parameters for function calls, unless stated otherwise.

### Read processing

Sequence reads were aligned to the *D. melanogaster* release 6 reference genome (dm6) using Bowtie (Langmead et al., 2009) version 1.1.2 (parameter –m 1) for ChIP-seq and DIP-seq samples and Bowtie2 (Langmead and Salzberg, 2012) version 2.2.9 (parameters -- very-sensitive, --no-discordant, --no-mixed, -X 100) for ATAC-seq samples considering read-pairs as non-nucleosome if *≤*100 bp according to (Buenrostro et al., 2013).

### Peak calling, robust peak sets and HAS definition

Peaks were called using Homer (Heinz et al., 2010) version 4.9.1 calling function findPeaks for DIP- and ChIP-seq samples using the corresponding input sample as control (parameters–style factor, -size 200, -fragLength 200, -inputFragLength 200 and –C 0) and for ATAC-seq samples without control (parameters -style dnase, -C 0, -gsize 137e6, -minDist 50). Peaks were defined as robust if the region was called in at least two replicate samples. HAS regions were used as defined by (Villa et al., 2016) with 309 HAS in total, of which 304 are located on the X chromosome and 5 on autosomes.

### X chromosomal enrichment

The X chromosomal enrichment was defined as the ratio of X chromosomal peak density over autosomal peak density and peak density was calculated as the number of peaks divided by chromosome length as defined by (Villa et al., 2016).

### *De novo* motif discovery and MRE definition

Enriched motifs in peak region were discovered using MEME (Bailey et al., 2009) version4.11.4 (parameters -dna, -mod zoops and -revcomp). For discovering the PionX motif, the list of PionX sites in (Villa et al., 2016) was used. As described in reference (Villa et al., 2016), we refer to MRE as the motif discovered within HAS, which is highly similar to the originally defined motif (Alekseyenko et al., 2008; Straub et al., 2008).

### Motif search

Motif search in peak region was performed with FIMO (Grant et al., 2011) version 4.11.4 (parameters --qv-thresh, --thresh 0.2, --max-stored-scores 1e6) applying a fifth-order background model.

### Browser profiles

Browser profiles were generated using tsTools (R) (https://rdrr.io/github/musikutiv/tsTools/) using mean per base read count per million mapped reads (rpm) of biologicals replicates, after aligned reads were extended to 200 bp fragments.

### Assignment of peak regions to chromatin states

Peak regions within one of the chromatin states were assigned to the corresponding chromatin state defined by (Kharchenko et al., 2011) and peaks overlapping with multiple chromatin states were assigned as ‘none’.

### ChIP-seq analysis

For calculating ChIP-seq signal enrichment as log_2_ ratio of IP over input, aligned reads were extended to 200 bp fragments and reads overlapping with target regions requiring a minimal overlap of half read length were counted and normalized to million mapped reads (rpm). For generating heat maps by calling the function pheatmap (R) and average profiles in a 2 kb window around HAS, coverage vectors were calculated as mean per base read count of IP samples and normalized to rpm. To test for statistical significant differences between *gst* RNAi and *trl* RNAi condition in CLAMP ChIP-seq, signal enrichment for the merged set of robust peak sets of both samples was calculated. Testing for difference was performed using limma (Ritchie et al., 2015) (R) including batch variables as random effect and calling the functions lmFit, eBayes (parameters trend = T, robust = T) and topTable (parameter adjust.method = ‘fdr’). Regions were defined as statistical significant different between conditions with FDR < 0.05.

### DIP-seq analysis

As proxy for MSL2-FLAG *in vitro* genomic binding we used the previously published DIP-seq experiment by (Villa et al., 2016) (GSE75033), where MSL2 was immunoprecipitated using *α*-FLAG M2 beads (Sigma-Aldrich) directed against the FLAG-tag, here referring to as ‘MSL2 (*α*-FLAG) – Villa’. For calculating DIP-seq signal enrichment as log_2_ ratio of IP over input, aligned reads were extended to 200 bp fragments and reads overlapping with target regions requiring a minimal overlap of half read length were counted and normalized to rpm. For generating heat map by calling the function pheatmap (R), euclidean distance between the enrichment for all biological replicates at combined robust peak set was measured by calling the function dist (R) and samples were hierarchical clustered using the ‘complete’ method by calling the function hclust (R). The row dendrogram was divided into 18 clusters by calling the function cutree (R) (parameter k = 18). For predicting the DNA roll between the base pairs at position +1 and +2 of the PionX motif, the best matching motif found by FIMO within the peak region was extended by 2 nucleotides at each end. The roll was predicted using DNAshapeR (Chiu et al., 2016) (R) by calling the function getShape. The DNA roll between the first two base pairs was assigned to the first base pair for simplicity, therefore referring to as ‘roll at position +1’. The number of GA:TC dinucleotides in the peak region was counted using rDNAse (R) by calling the function kmer considering also the reverse complement. We calculated the GA:TC dinucleotide density as the number of GA:TC-dinucleotides divided by the peak length. The length of GA:TC repeats was counted using Biostrings (R) by calling the function vcountPattern considering also the reverse complement.

### ATAC-seq analysis

For calculating ATAC-seq signal intensities, fragments on the + strand were moved by +4 bp and on the - strand by −5 bp according to (Buenrostro et al., 2013). Signal intensities in target regions were calculated by calling summarizeOverlaps (parameter ignore.strand = TRUE) from GenomicAlignments (Lawrence et al., 2013) (R). Further analysis was performed using DESeq2 (Love et al., 2014) (R), including batch variables as random effect. Sites were considered as statistical significant different between conditions with absolute log_2_ fold-change > 0.5 and FDR < 0.1 by calling the function results (parameters lfcThreshold = 0.5, altHypothesis = “greaterAbs”). For analyzing ATAC-seq signal at HAS, the size factor obtained by using the combined robust peak sets of the corresponding samples was used. For comparing ATAC-seq signals in Kc cells with S2 cells, only X chromosomal peak regions were considered.

## Acknowledgements

We thank S. Krebs and the LAFUGA Genomics Facility for next generation sequencing,M. Müller for initial cloning of full-length CLAMP-FLAG, E. Kremmer for antibody generation, and E. Larschan for the antibody against CLAMP. This work was supported by the Deutsche Forschungsgemeinschaft through grant Be1140/8-1 to PBB. OM was supported by RSF #17-74-20155. CA acknowledges a DFG fellowship from the Graduate School for Quantitative Biosciences Munich (QBM).

## Author contributions

CA, CR, and PBB conceived the study; CA performed all experiments except for co-immunoprecipitation and Yeast-Two-Hybrid experiments; SK expressed proteins and performed co-immunoprecipitation experiments; ET and OM conceived, performed and analyzed Yeast Two-Hybrid experiments; CA performed bioinformatics analyses; All authors analyzed data; CR and PBB provided feedback and supervision; CA, CR, and PBB wrote the manuscript; PBB secured funding.

## Expended View Figures

**Figure EV 1.**
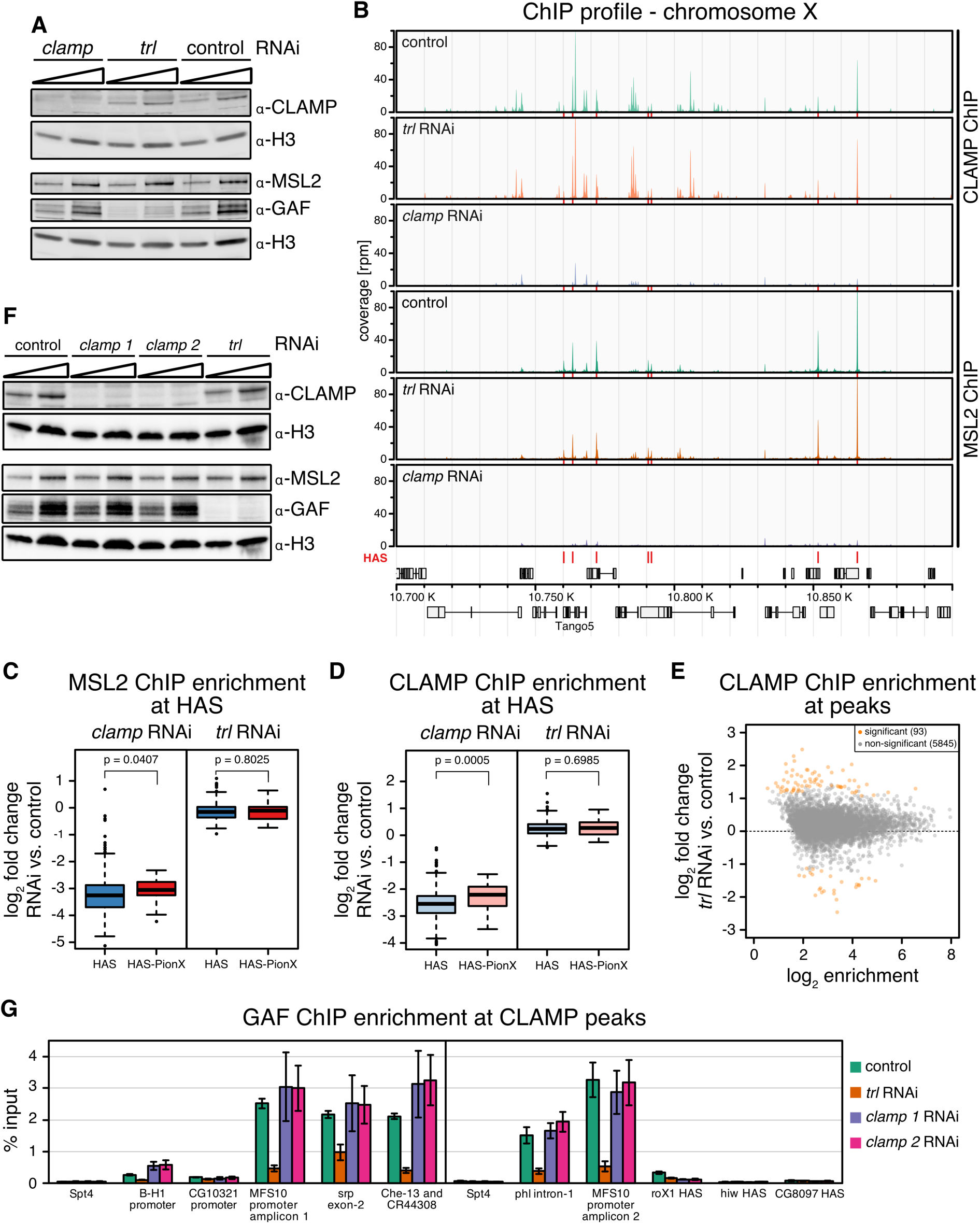
Non-competitive genomic binding of CLAMP and GAF *in vivo*. **(A)** Western blot detection of MSL2, CLAMP, GAF and H3 of whole cell extract from control S2 cells and after *clamp* or *trl* RNAi, which were used for ChIP-seq. **(B)** Genome browser profile of CLAMP ChIP seq (upper panels) and MSL2 ChIP seq (lower panels) profiles as indicated to the right showing mean coverages (control and *trl* RNAi: n = 3, *clamp* RNAi: n = 4) along a representative 200 kb window on the X chromosome as indicated. Red bars above the gene models indicate position of HAS. **(C)** Boxplot of mean log_2_ fold-change of MSL2 ChIP-seq enrichment after *clamp* RNAi (left panel) and *trl* RNAi (right panel) versus control at 309 HAS grouped as HAS non-overlapping with PionX sites (HAS, blue, n = 272) and overlapping with PionX sites (HAS-PionX, red, n = 37). Unpaired Mann-Whitney-test was used to test for difference between groups, p-values are shown. **(D)** Boxplot of mean log_2_ fold-change of CLAMP ChIP-seq enrichment after *clamp* RNAi (left panel) and *trl* RNAi (right panel) analogous to [C]. **(E)** Scatter plot of mean log_2_ fold-change (n = 3) of CLAMP ChIP enrichment in S2 cells after *trl* RNAi versus control against mean log_2_ read count in control t at robust CLAMP peaks (n = 5938). Sites with statistically significant different CLAMP enrichment (|lfc| > 0 and fdr < 0.05) are marked in orange (n = 93). **(F)** Western blot detection of MSL2, CLAMP, GAF and H3 in whole cell extracts from control S2 cells and after *clamp* (2 different dsRNA constructs) or *trl* RNAi, which were used for ChIP-qPCR. **(G)** Bar chart of GAF ChIP enrichment relative to input in control S2 cells and after *clamp* (2 different dsRNA constructs) or *trl* RNAi, determined by qPCR at nine loci overlapping with robust CLAMP ChIP-seq peaks: three HAS occupied by MSL2 and CLAMP but not GAF in control S2 cells (*roX1* HAS, *hiw* HAS-PionX, and *CG8097* HAS-PionX), two sites occupied by CLAMP but not GAF (*B-H1* promoter and *CG10321* promoter), four sites which are bound by CLAMP and GAF (*MFS10* promoter, *srp* exon-2, *phl* intorn-1 and ‘between *Che-13* and *CG44308*’) and the *spt4* locus as negative control. Error bars show the standard error of the mean from 4 biological replicates, each representing the mean of technical triplicates.

**Figure EV 2.**
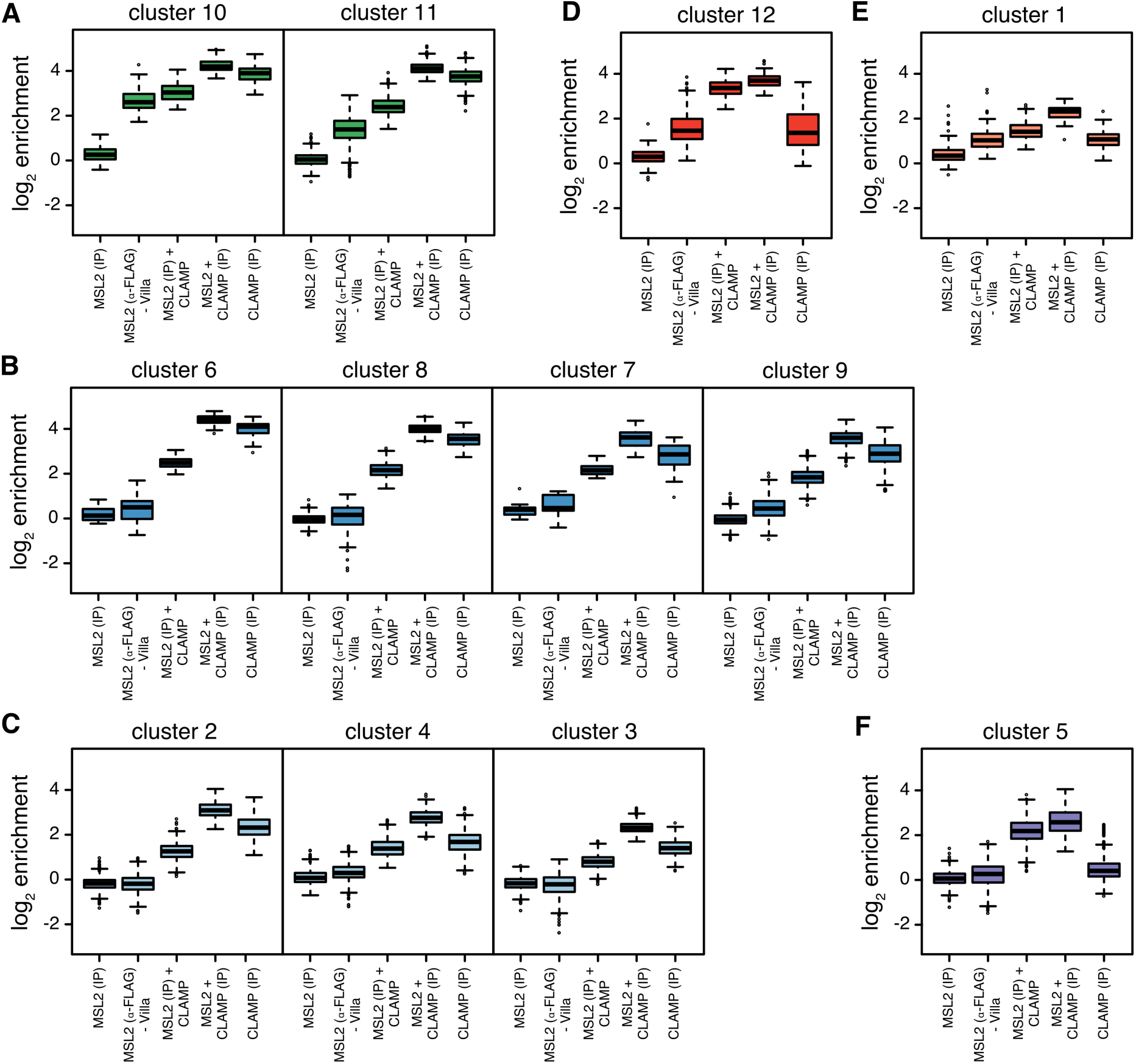
DNA binding cooperativity between of MSL2 and CLAMP *in vitro*. **(A)** Boxplot of mean log_2_ enrichment (n = 3) of *in vitro* DIP-seq at peaks belonging to the clusters 10 and 11 as defined in (Figure 4A). The categories are indicated: MSL2 with *α*-FLAG IP from(Villa et al., 2016) as proxy for MSL2 *in vitro* binding, MSL2 with *α*-MSL2 IP, MSL2 and CLAMP with *α*-MSL2 IP, CLAMP with *α*-CLAMP IP, and MSL2 and CLAMP with *α*-CLAMP IP. **(B)** For clusters 6 to 9 as in (A). **(C)** For clusters 2 to 4 as in (A). **(D)** For cluster 12 as in (A). **(E)** For cluster 1 as in (A). **(F)** For cluster 5 as in (A).

**Figure EV 3.**
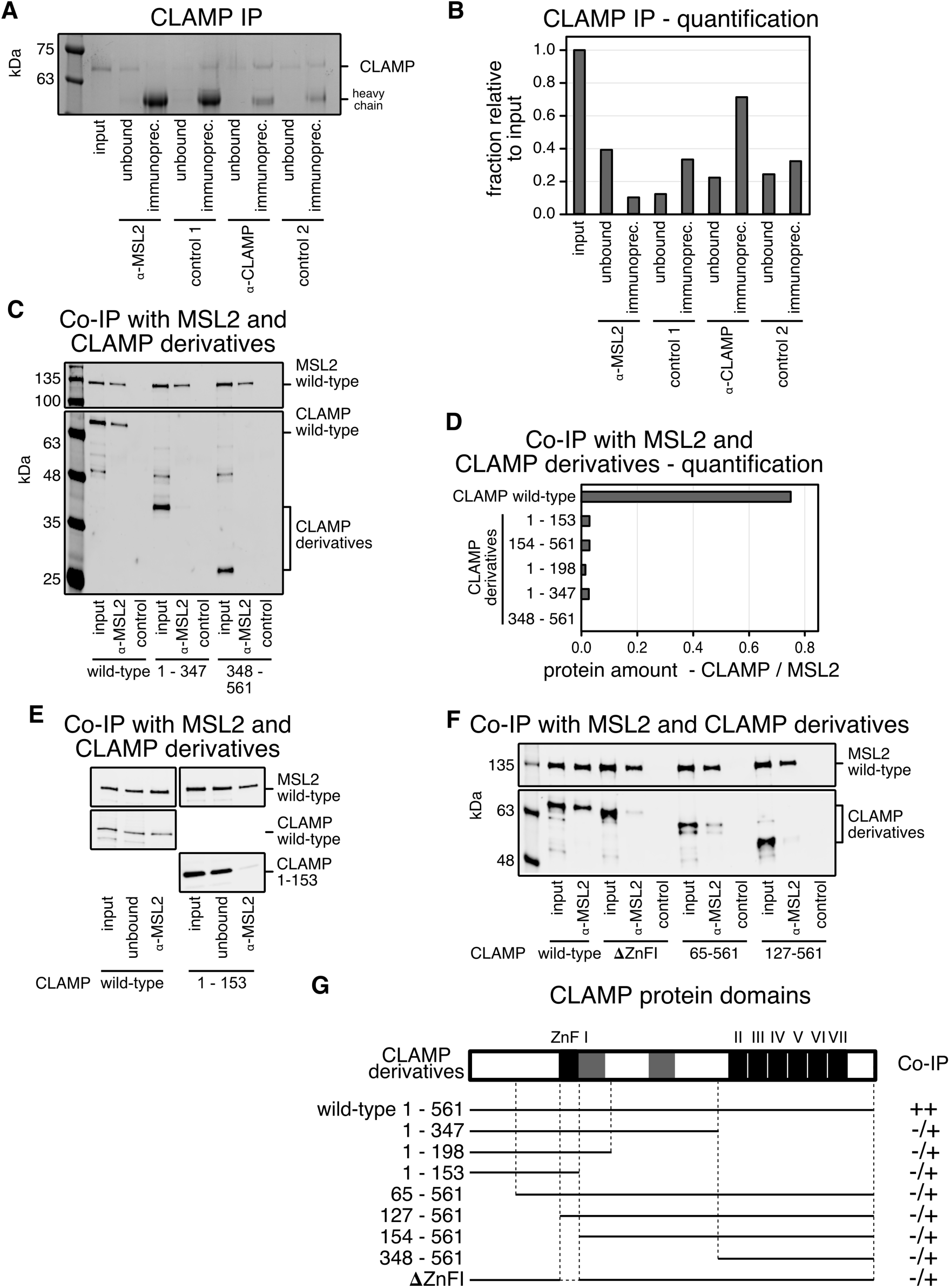
CLAMP and MSL2 form a stable complex. **(A)** SDS-PAGE analysis with Coomassie staining of IP fractions. Purified wild-type recombinant CLAMP-FLAG was immunoprecipitated with *α*-MSL2 serum and the corresponding pre-immune serum as control 1, and with specific affinity purified *α*-CLAMP antibody mixed into an irrelevant rabbit serum and with the irrelevant rabbit serum only as control 2. The corresponding unbound fractions are loaded next to each IP. Molecular weight markers are shown on the left. **(B)** Bar chart of the quantification from IP experiments shown in (A). The protein amount in the unbound fractions and IP’s were quantified relative to the input. **(C)** Quantitative Western blot analysis using *α*-FLAG antibody of co-IP experiments with extracts from Sf21 cells expressing wild-type MSL2-FLAG and CLAMP-FLAG deletion mutants. Co-IP was performed with *α*-MSL2 serum and the corresponding pre-immune serum as control 1. IP fractions were loaded next to each corresponding input. **(D)** Bar chart of the ratio between CLAMP and MSL2 in co-IP experiments as shown in (C). **(E)** Quantitative Western blot analysis using *α*-FLAG antibody of co-IP experiments with extract from Sf21 cells expressing wild-type MSL2-FLAG and 3-times more concentrated extract expressing wild-type CLAMP-FLAG and N-terminal CLAMP^1-153^ fragment. Co-IP was performed with *α*-MSL2 serum. **(F)** Quantitative Western blot analysis using *α*-FLAG antibody of co-IP experiments as in (E) for CLAMP derivatives with various deletions in the N-terminal half: CLAMP^ΔZnFI^, CLAMP^65-561^ and CLAMP^127-561^. **(G)** Summary of MSL2 and CLAMP interaction from co-IP experiments as presented in (C). Interaction with full length CLAMP is depicted by (++) and interactions above background by (-/+). CLAMP domain architecture is drawn to scale. White rectangles represent the conserved regions (Appendix S8). Black rectangles represent the seven Zinc-finger domains. The two grey boxes represent the prion-like domains according to (http://plaac.wi.mit.edu) (Kaye et al., 2018).

**Figure EV 4.**
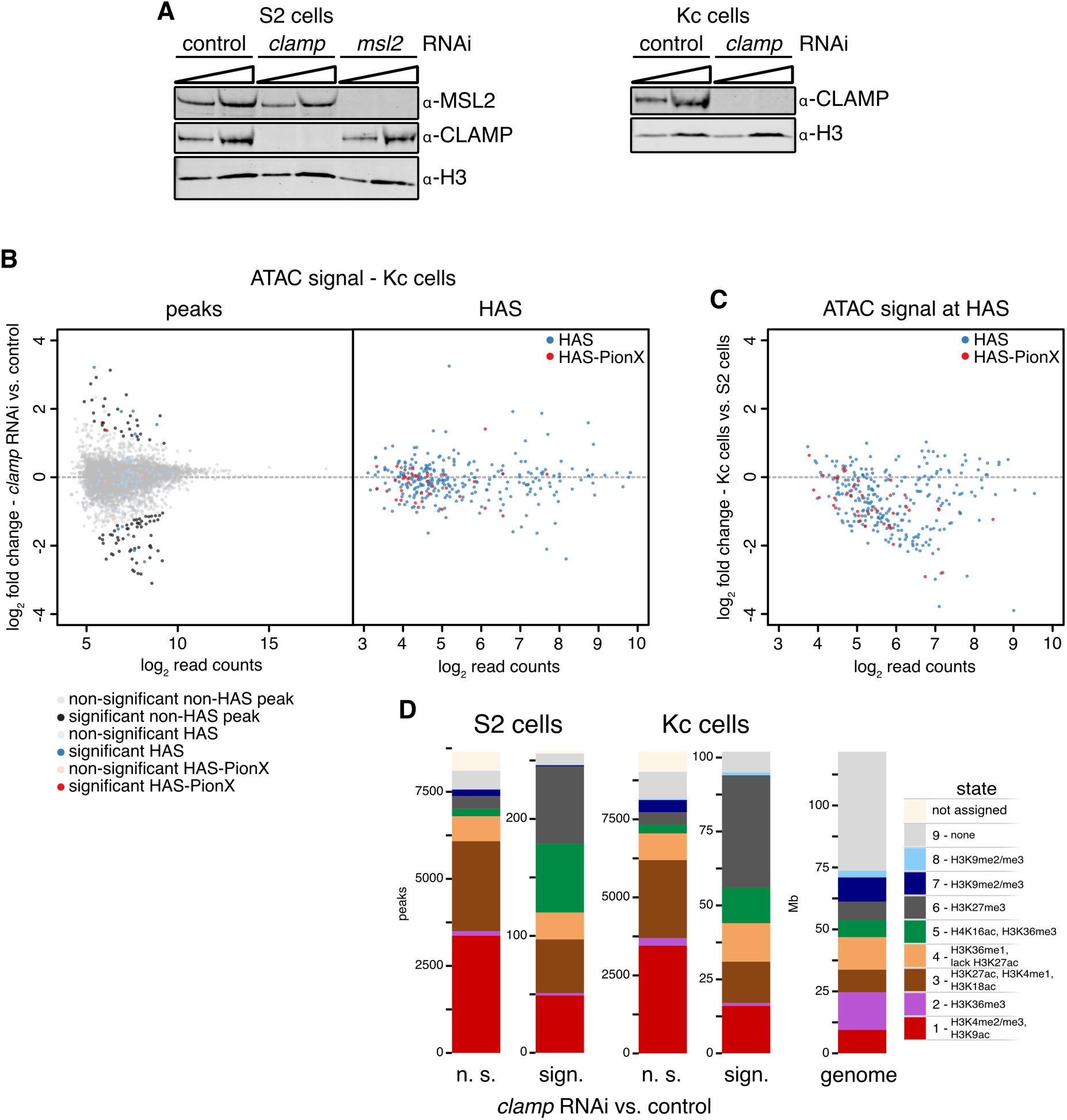
Most HAS are inaccessible in females independently of CLAMP. **(A)** Western blot of CLAMP, H3 and MSL2 (only for S2 cells) of whole cell extract from control S2 cells and after *clamp* and *msl2* RNAi (left panel) and from control Kc cells and after *clamp* RNAi (right panel). These cells were used for ATAC-seq. **(B)** Scatter plot of mean log_2_ fold-change (n = 3) of ATAC-seq signal in Kc cells upon *clamp* RNAi versus control against mean log_2_ read count in control at robust ATAC peaks (left penal, n = 9767) and at HAS (right penal, n = 309). Left panel, HAS non-overlapping with PionX sites (HAS) are marked in blue and overlapping with PionX sites (HAS-PionX) are marked in red (the remaining sites are displayed in grey). Sites with statistically significant different ATAC signal between *clamp* RNAi and control condition (|lfc| > 0.5 and fdr < 0.1) are marked in darker color. 102 ATAC peaks are statistically significant different between conditions, including 10 HAS and 1 HAS-PionX. Right panel, HAS non-overlapping with PionX sites (HAS, n = 272) are marked in blue and overlapping with PionX sites (HAS-PionX, n = 37) are marked in red. **(C)** Scatter plot of mean log_2_ fold-change (n = 4) of ATAC-seq signal in Kc cells versus S2 cells against mean log_2_ read count in S2 cells at HAS (n = 309). HAS non-overlapping with PionX sites (HAS, n = 272) are marked in blue and overlapping with PionX sites (HAS-PionX, n = 37) are marked in red. **(D)** Bar chart of chromatin state distribution of robust ATAC peaks in S2 cells (left panels, n = 8913) grouped as not statistical or statistical significantly different between *clamp* RNAi and control (|lfc| > 0.5 and fdr < 0.1) as defined in (Figure 7B) and for Kc cells (middle panels, n = 9767) as defined in (B) and cumulative chromatin state sizes (right penal, genome) as reference for uniform distribution.

## Appendix

### Appendix Figure Legends

**Appendix S1. CLAMP peaks are located mainly at promotors and enhancers.**

Bar chart of chromatin state distribution of robust CLAMP peaks in S2 cells (n = 5938) grouped as not statistical or statistical significantly different between *trl* RNAi and control conditions (|lfc| > 0 and fdr < 0.05) as defined in (Figure EV1E) and cumulative chromatin state sizes (genome) as reference for uniform distribution.

**Appendix S2. MSL2-FLAG and CLAMP-FLAG preparation.**

SDS-PAGE with Coomassie staining of three biological independent CLAMP-FLAG and MSL2-FLAG purification used for DIP-seq and co-IP experiments. Next to each protein purification are BSA standards loaded for quantification. Molecular weight markers are shown to the left.

**Appendix S3. Distinct sequence feature sets determine MSL2 and CLAMP binding *in vitro*.**

Clustered heat map of *in vitro* DIP-seq signal enrichment (as shown in Figure 4). Score of the best matching MRE motif, score of the best matching PionX motif, the roll at position 1 of the best matching PionX motif, the density of GA:TC dinucleotides and the length of GA:TC repeats are indicated at the right. *De novo* discovered motif within each cluster are shown to the right.

**Appendix S4. Sequence features of clusters of genomic *in vitro* binding sites for MSL2 and CLAMP.**

**(A)** Summary of the peaks in each clusters of the heatmap in (Fig. 4).

**(B)** Boxplot of peak features grouped into to the twelve clusters defined in Figure 4. Panels from left to right show: Score of the best matching MRE motif, score of the best matching PionX motif, the roll at position 1 of the best matching PionX motif, the density of GA:TC dinucleotides and the length of GA:TC repeats.

**(C)** Bar chart of X chromosomal enrichment of peaks grouped into the clusters described in (Fig. 4).

**Appendix S5. Protein-interaction of CLAMP and MSL2 in yeast 2-hybrid assays.**

**(A)** Yeast 2-hybrid (Y2H) test of CLAMP and MSL2 interaction. The activating domain of Gal4 (AD) was fused to CLAMP and the DNA binding domain of Gal4 (DBD) to MSL2. Upper panel show representative Y2H assays. Lower panel summarizes all tested fragments. Numbering correspond to the amino acids included in the constructs. White boxes depict the conserved regions in CLAMP and MSL2. Black boxes indicate known domains and grey boxes indicate Prion-Like domains. The CLAMP-PB isoform was used in Y2H (1-566 amino acids) while CLAMP-PA was used for co-IP. The ‘+’ and ‘-’ signs refer to the presence and absence of interaction, respectively.

**(B)** Alignment of the CXC-containing C-termini from MSL2 homologs from 12 *Drosophila* species (Dmel: *D. melanogaster* MSL2-PA 507-773; Dsec: *D. sechellia* GM11132-PA 506-775; Dsim: *D. simulans* GD15895-PB 514-783; Dyak: *D. yakuba* GE14867-PA 513-780; Dere: *D. Erecta* GG24438-PA 513-780; Dana: *D. ananassae* GF23410-PA and PB 433-723; Dpse: *D. pseudoobscura* GA16882-PA/PB 771-1081/792-1102, respectively; Dper: *D. persimilis* GL18894-PA 792-1100; Dvir: *D. virilis* MSL2-PA and PB 495-760; Dmoj: *D. mojavensis* GI14252-PA/PB/PC 591-85/591-858/591-857, respectively; Dgri: *D. grimshawi* GH10188-PB 546-821; Dwil: *D. willistoni* GK23860-PB 735-977) using MultAlin (Corpet, 1988). The homologs from *D. mojavensis* and *D. virilis* have extended divergent C-termini, which have been truncated for clarity of the alignment (*). In the CXC domain (47 AA), 40% of the residues are conserved to more than 90% (in red) and 30% of the residues are 50-90% conserved (in blue). In the conserved region 2 (CR2) ranging from AA 620-685, 32% of AA are more than 90% conserved and 50% of AA are 50-90% conserved. The C-terminal CR3 (28 AA) is conserved from 50-100% on 86% of its length.

**Appendix S6. All MSL2 deletion constructs containing the RING domain interact with MSL1 in yeast 2-hybrid assays.**

**(A)** Replicates including controls for yeast 2-hybrid (Y2H) screening of CLAMP and MSL2 interaction. Full-length or 1-153 amino acid fragment of CLAMP was fused to the GAL4 activating domain (AD) and tested for interaction with MSL2 fused to the GAL4 DNA-binding domain (DBD). CLAMP fusions were tested for the absence of interaction with the GAL4 DNA-binding domain alone (pGBT). MSL2 was tested for the absence of interaction with the GAL4 activating domain alone (pGAD).

**(B)** MSL2 binds MSL1 in Y2H assay. Different fragments of MSL2 were fused to the GAL4 activating domain (AD) and tested for interaction with MSL1 fused to the GAL4 DNA-binding domain. MSL2 fusions were tested for the absence of interaction with the GAL4 DNA-binding domain (DBD) alone. Numbering correspond to the amino acids included in the constructs. White boxes depict the conserved regions in MSL2 and black boxes indicate known domains. The ‘+’ and ‘-’ signs refer to the presence and absence of interaction, respectively.

**Appendix S7. CLAMP is conserved over its entire length within *Drosophila* species.**

Alignment of CLAMP homologs from 11 *Drosophila* species (Dmel: *D. melanogaster* CG1832-PA; Dsec: *D. sechellia* GM16114-PA; Dmel: *D. melanogaster* CG1832-PB; Dyak: *D. yakuba* GE12967-PA; Dere: *D. Erecta* GG21465-PA; Dpse: *D. pseudoobscura* GA14887-PA; Dper: *D. persimilis* GL26169-PA; Dana: *D. ananassae* GF13861-PA; Dgri: *D. grimshawi* GH10979-PA; Dvir: *D. virilis* GJ15143-PA; Dmoj: *D. mojavensis* GI17493-PA; Dwil: *D. willistoni* GK15349-PA) using MultAlin (Corpet, 1988).

**Appendix S8. Conservation of MSL2 sequence between 12 *Drosophila* species.**

Alignment of MSL2 homologs from 12 *Drosophila* species as in Appendix S5B.

### Appendix Table Legends

**Appendix Table S1. Generation of constructs for Yeast two-hybrid assay.**

Primer sequences used for cloning of CLAMP and MSL2 derivatives.

**Appendix Table S2. Generation of constructs for interaction domain mapping.**

Primer sequences used for cloning of CLAMP and MSL2 derivatives.

**Appendix Table S3. Amplicons for ChIP-qPCR.**

Primer sequences used qPCR after ChIP.

---

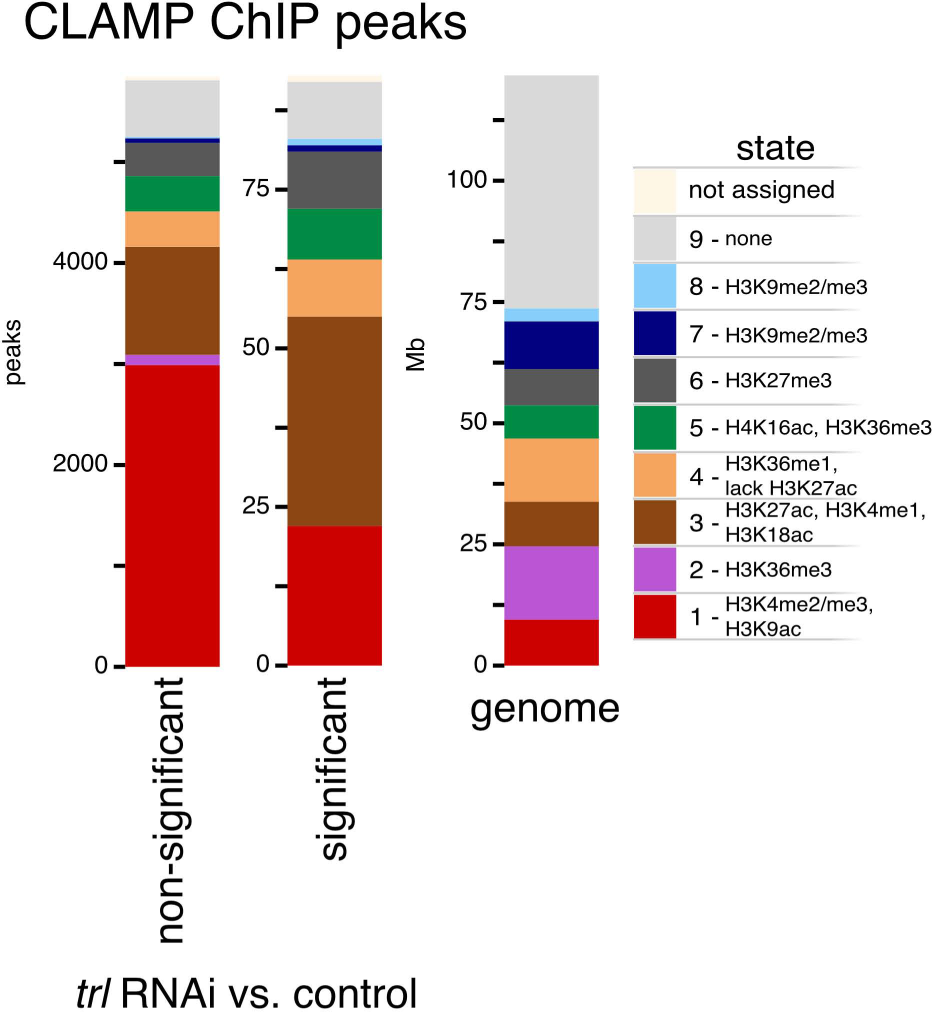

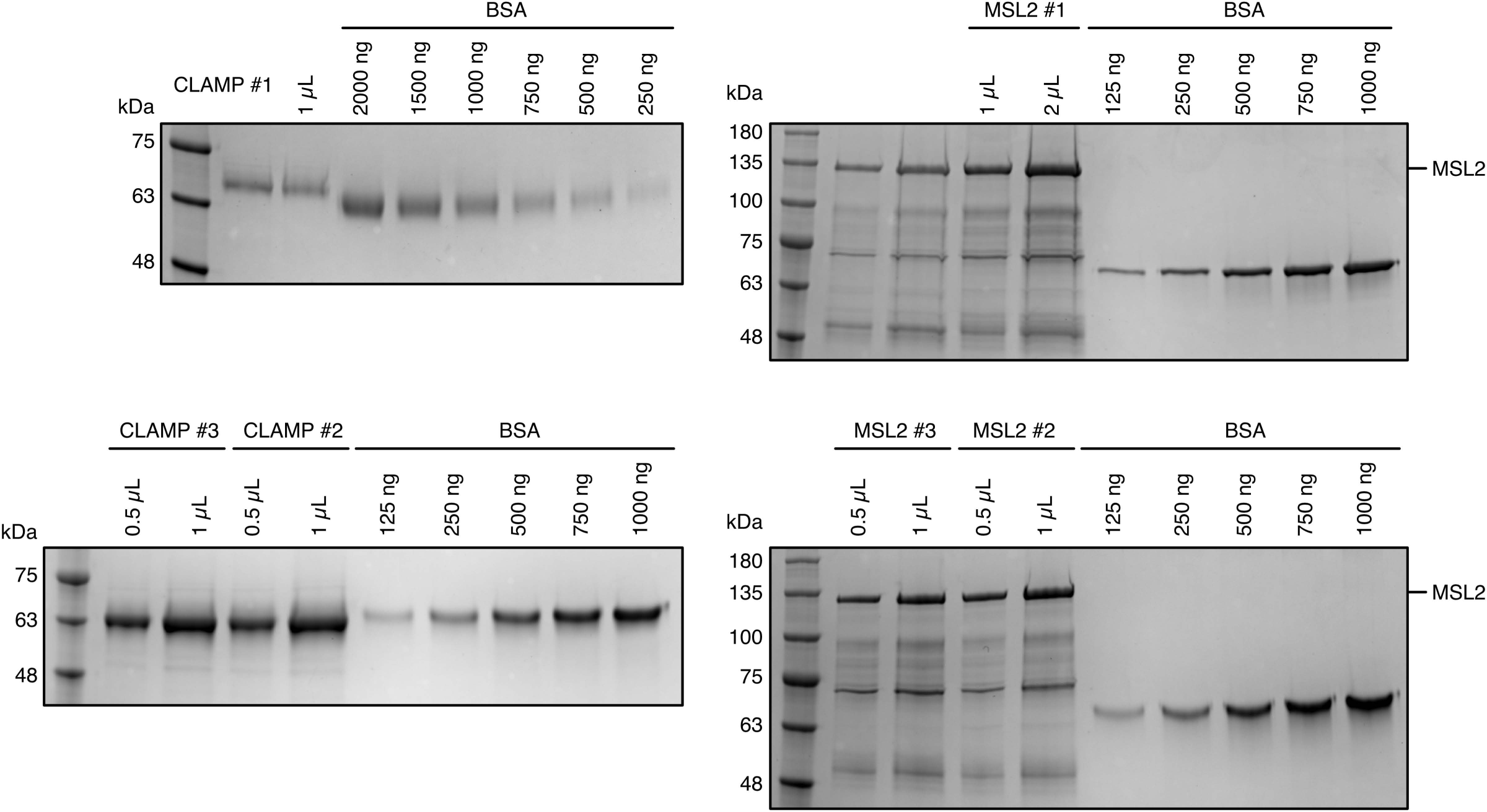

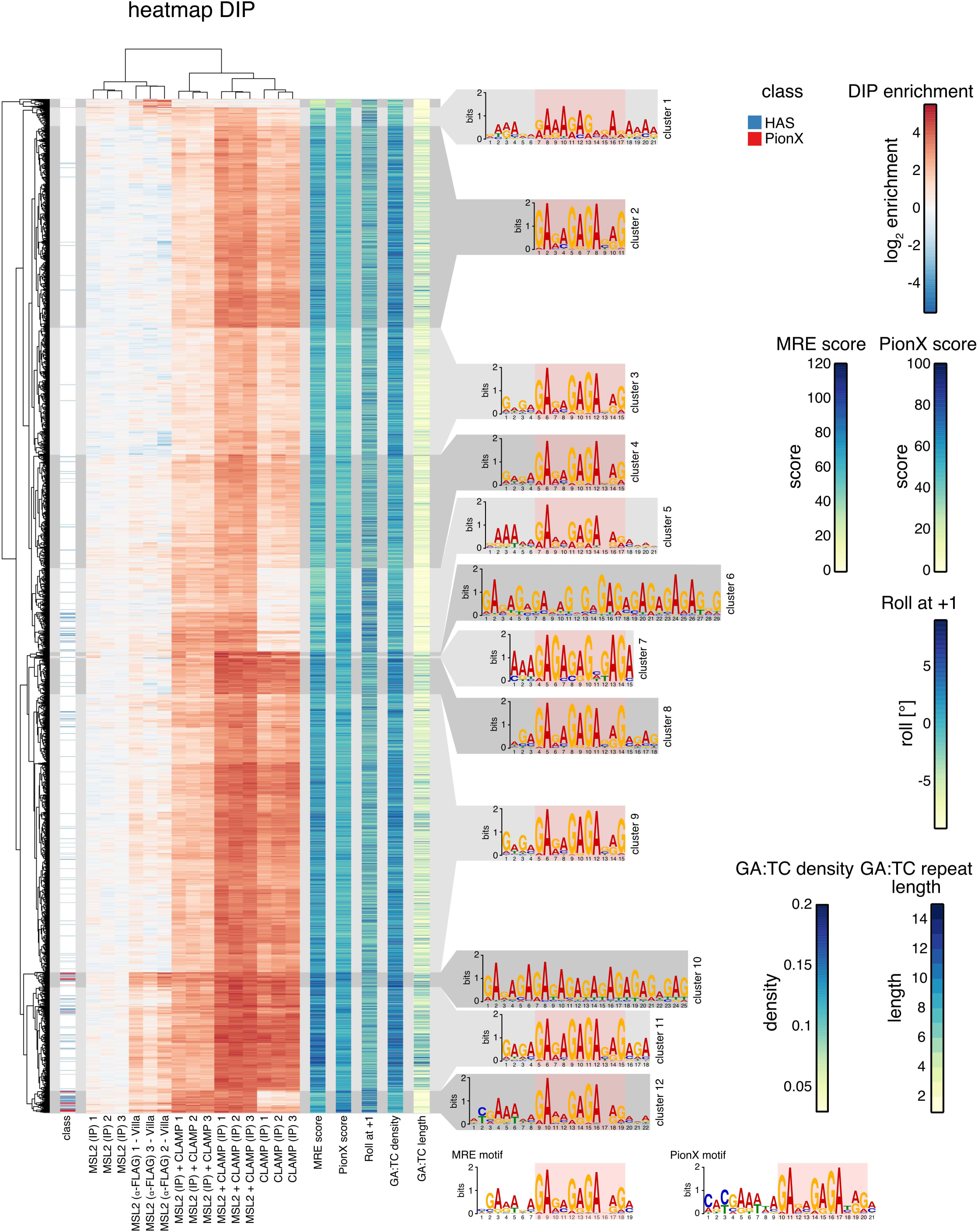

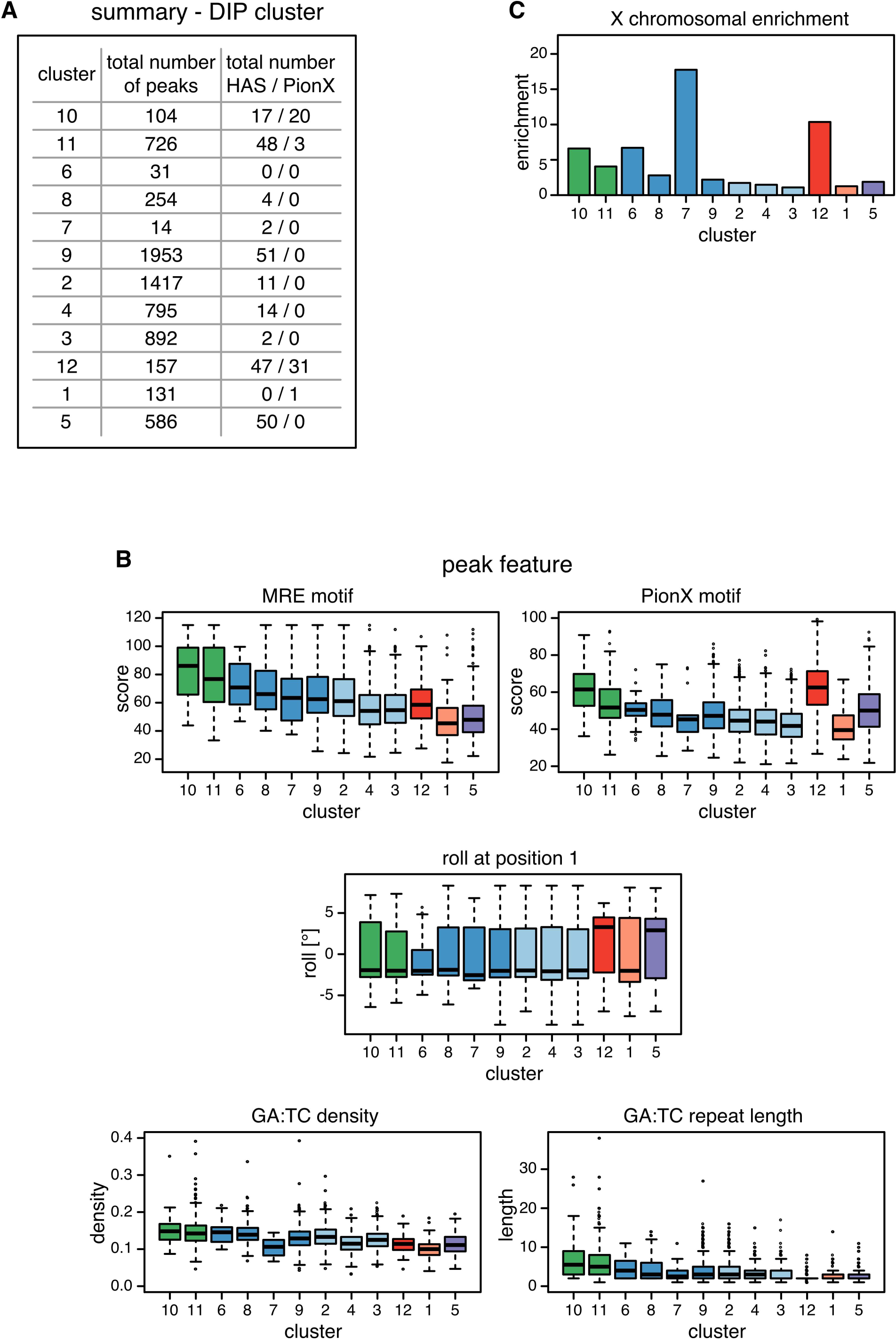

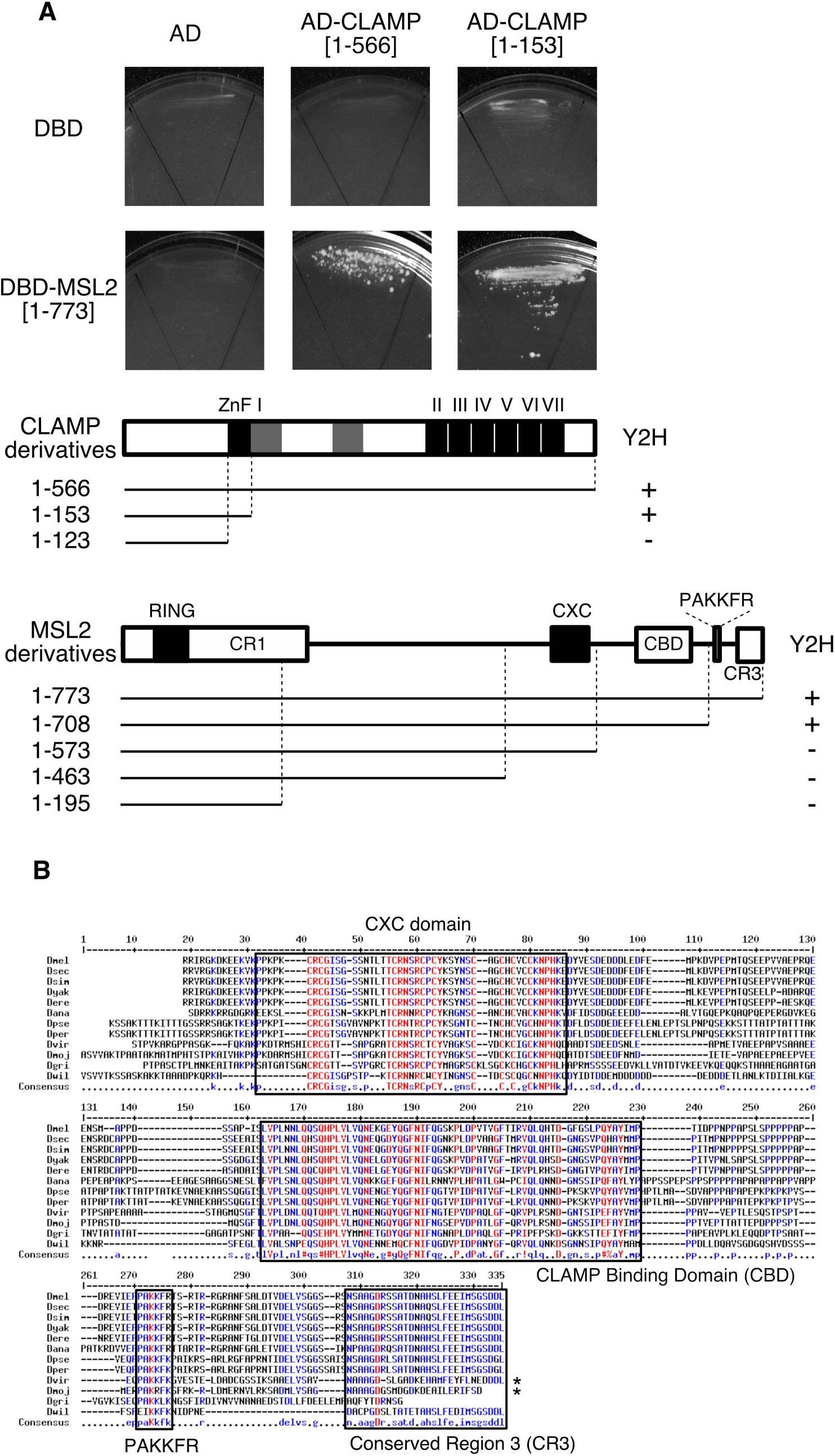

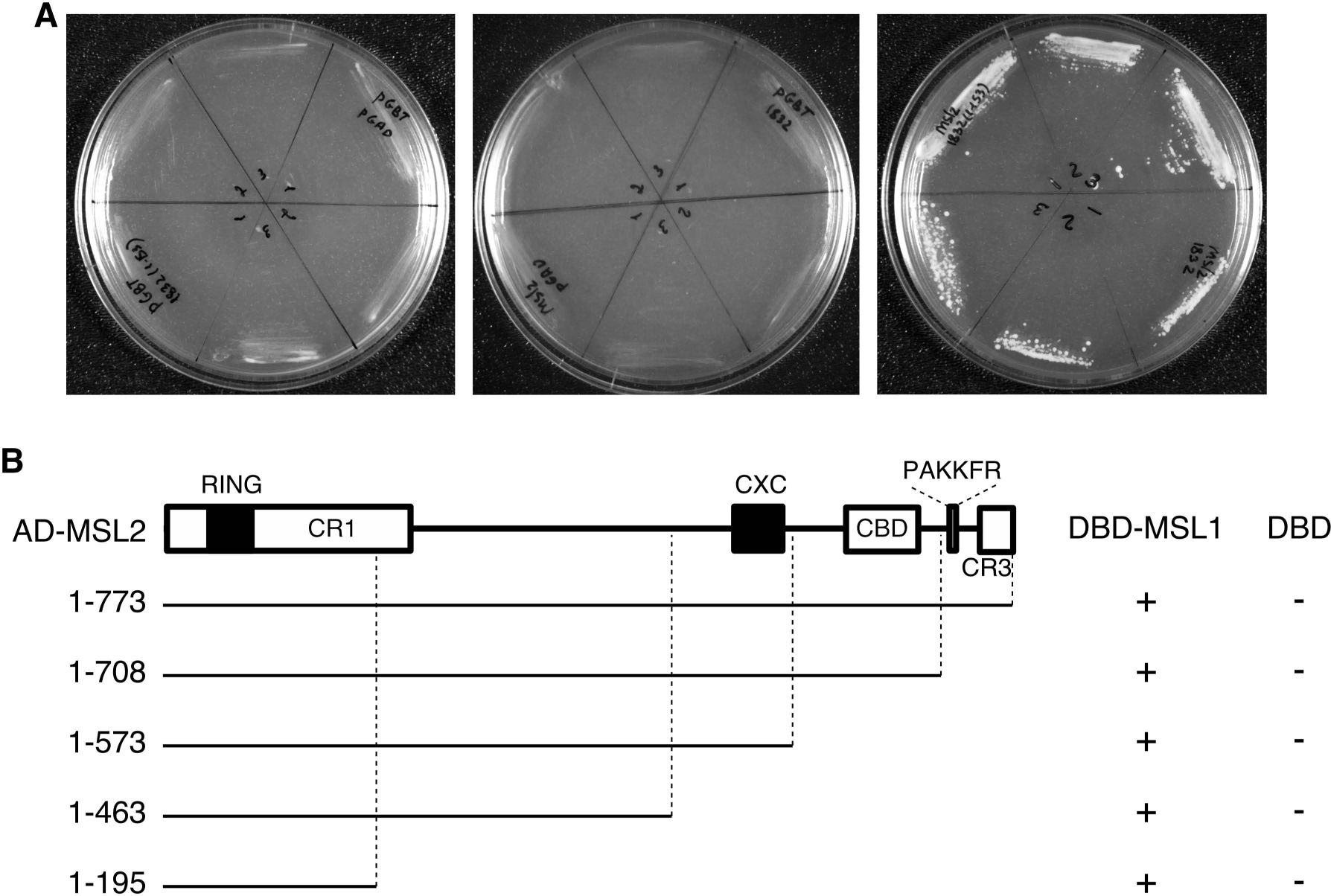

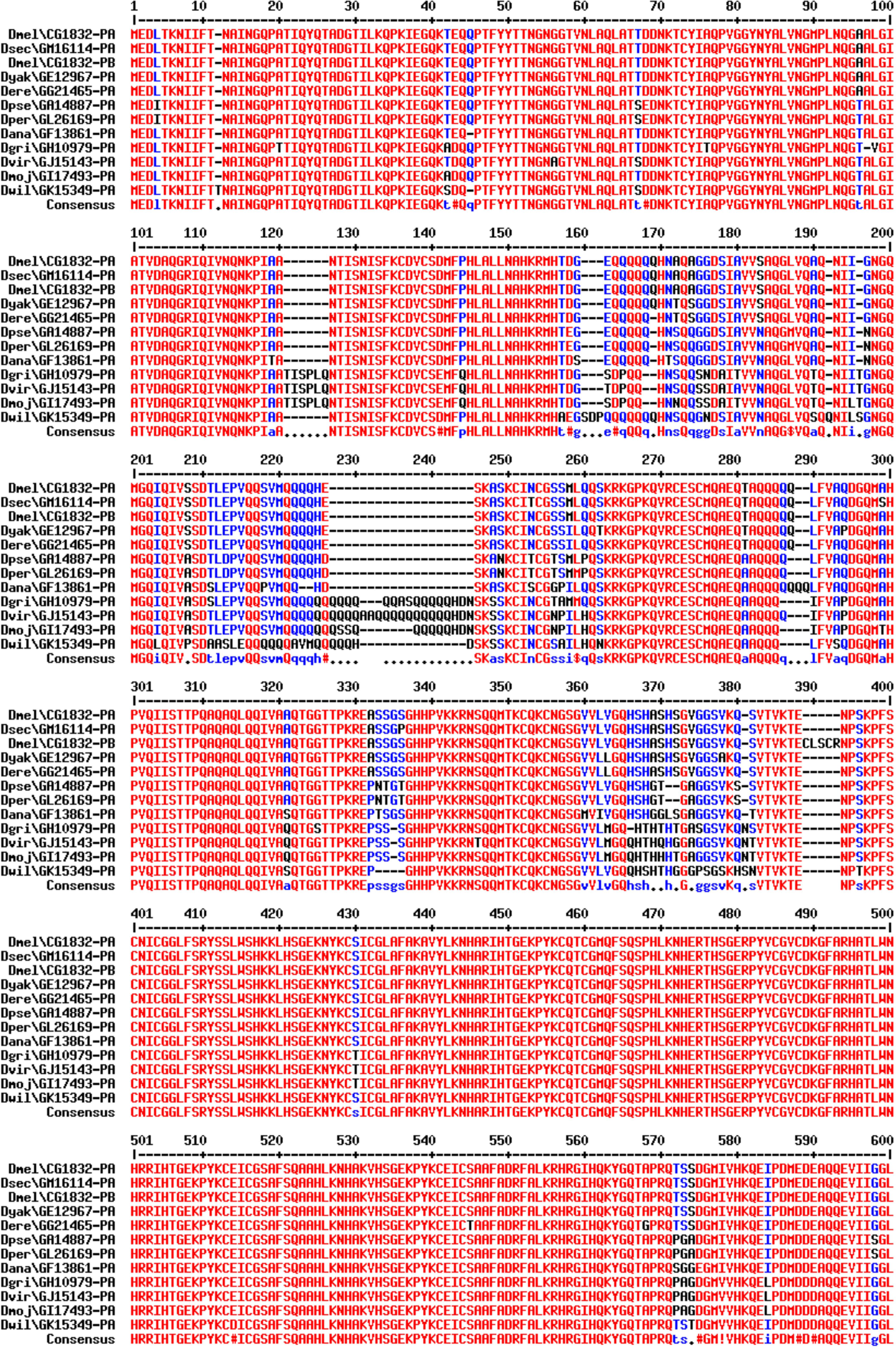

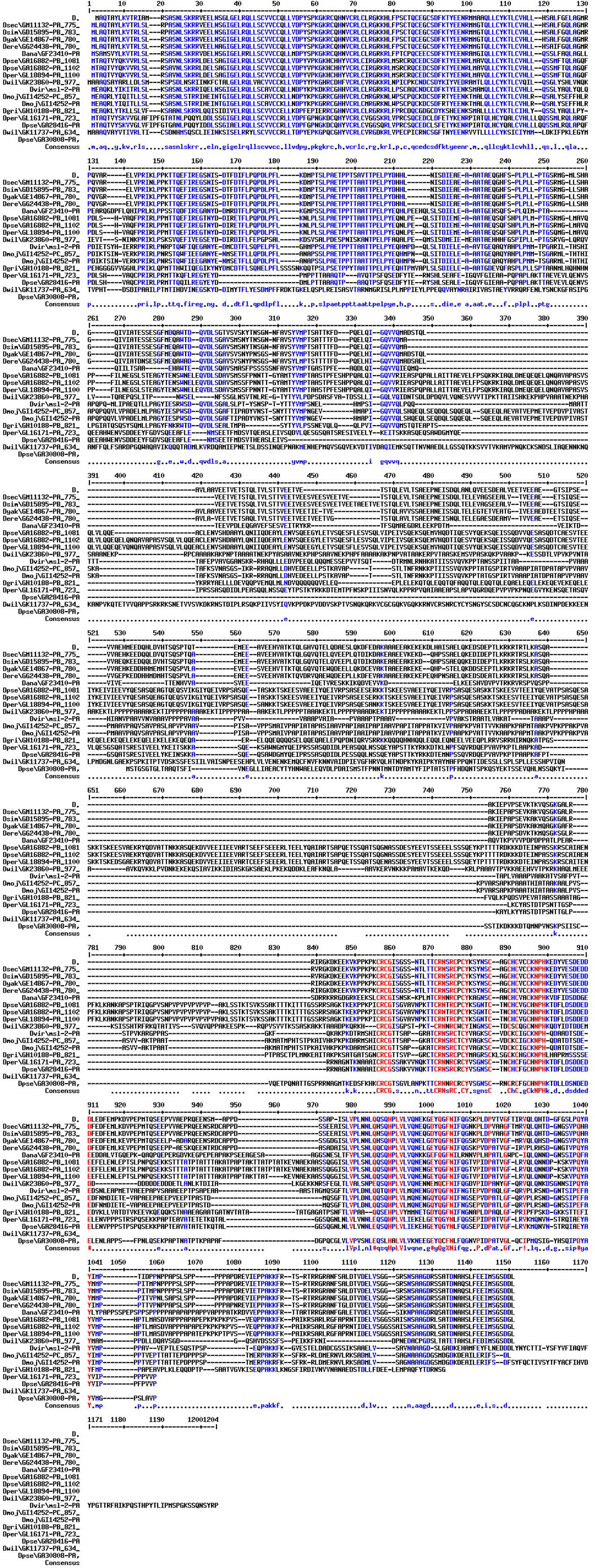

